# Cytocapsular cancer evolution analyses of 311 kinds of cancers

**DOI:** 10.1101/2022.07.08.499362

**Authors:** Tingfang Yi, Gerhard Wagner

**Affiliations:** Cytocapsula Research Institute, 245 First Street, Cambridge, MA, 02142 USA; Cellmig Biolabs Inc., 245 First Street, Cambridge, MA 02142 USA; Department of Biological Chemistry and Molecular Pharmacology of Harvard Medical School, 240 Longwood, Boston, MA, 02115 USA

## Abstract

Cancer is a leading cause of human lethality. Cytocapsular tube, a newly discovered cancer cell specific organelle *in vivo*, plays pleiotropic biological functions and its generation distinguishes incomplete from complete cancer cells. It is essential for complete malignant tumor growth, cancer metastasis, conventional cancer drug pan-resistance, and cancer relapse. However, mechanisms of cytocapsular cancer evolution are still elusive. Here, we investigated cytocapsular cancer evolution in 311 kinds/subtype of cancers, including 265 types/subtypes of solid cancers and 46 types/subtypes of liquid/hematological cancers. We analyzed 9,856 pieces of annotated clinical tissue samples from 9,682 cancer patients in the asymptotic early stage, Stages I-IV, and before, during and after cancer treatments. We discovered that cytocapsular cancer evolution in **solid cancers** includes: transformation, formation of incomplete cancer cells, transition to complete cancer cells surrounded by cytocapsulae (CC), merging of complete cancer cells by devolution, formation of cytocapsular tubes (CCTs) and complete malignant tumors in superlarge CC. This is followed by generation of CCT networks, cancer metastasis, CCT network-tumor system (CNTS), CCT degradation and decomposition, and spatiotemporal moving CNTS. In addition, cytocapsular cancer evolution related to **liquid** (**hematological) cancers** including bone marrow, thymus, lymph nodes, and spleen, mirrors the process in solid cancers, except that cancer cells in the blood only form CCs but not CCTs. In summary, our study established a cytocapsular cancer evolution atlas, which may pave an avenue for the research on therapy of both solid and liquid cancers.

## Introduction

Cancer is the leading source of mankind lethality^1–4^. The correct understanding of the nature of cancer evolution is essential for cancer screening, diagnosis, prognosis and therapy. In 1859, Charles Darwin provided an evolutionary framework with three key conceptions of variation, heredity, and selection for the understanding of somatic selection, diversification and extinction^5^. In 1976, Peter Nowell hypothesized that tumorigenesis is also an evolutionary process, and Darwin’s principles could be applied to elucidate the underlying mechanisms of cancer formation and development^6^. In the last decades, genomic, metabolic, cellular, cellular-plastic, and sub-clonal aspects of cancer evolution have been intensively studied. Several models on tumor evolution have been hypothesized, including linear evolution, branched evolution, macroevolution, and neutral evolution, which are determined by diverse scales of evolution-driving factors: genetic alterations, genomic aberrations (discordant inheritance, DNA macro-alterations), cell plasticity, and microenvironments^6^. Cancer cell-specific addicted and essential genes and target proteins are selected for cancer drug discovery and therapy, including small molecule drugs, peptide drugs, miRNA drugs, antibody drugs, and immune drugs (reagents and immune cells), and all kinds of physical therapies. Tumors’ biochemical, biophysical and metabolic features are targeted for the invention of all kinds of cancer screening and diagnosis methods. Despite these huge efforts, about 10 million cancer deaths and around 20 million new cancer cases occur in 2020 alone worldwide^1^. Thus, the marginal cancer pharmacotherapy outcomes suggest that the nature of cancer evolution is still beyond our conventional understanding^7–13^.

In January, 2018, we reported the discovery of the cytocapsular tube (CCT), which functions as cancer cell translocation physical pathways^14^. From 2018 to 2021, we found and reported that the CCT is a cancer cell specific organelle in cancer patients *in vivo*: CCTs and CCT networks are associate with all kinds of clinical solid cancers in all stages and provide membrane-enclosed physical freeways for cancer metastasis, but do not appear in normal human tissues or clinical benign tumors^15–16^. In 2020, the first anti-TNSDF antibody recognizing CCT marker protein CM-01 (code, not real protein name) was approved and registered by USA FDA as Class I MVU for clinical application as complementary cancer diagnosis information. From 2021 to 2022, we found and reported pleiotropic biological functions and roles of CCs and CCTs in complete cancer cell generation, cancer metastasis, complete malignant tumor formation and growth, conventional cancer drug pan-resistance, cancer relapse, cancer evolution, and normal tissue/organ biological functional failure, and precise cancer diagnosis^17–19^.

Here, we investigated the cytocapsular cancer evolution of 311 kinds of solid and liquid cancers in 35 kinds of tissues and organs: adrenal gland, bladder, blood in blood vessel, blood in bone marrow, bone, bone marrow, brain, breast, cervix uteri, colon, esophagus, fibrous, gallbladder, head/neck, intestine, kidney, liver, lung, lymph node, oral cavity, ovary, pancreas, penis, prostate, rectum, skin, smooth muscle, spleen, stomach, testis, thymus, thyroid, uterus, and vulva. Our study unveiled a cytocapsular cancer evolution atlas in solid and liquid cancers, which will facilitate the research of cancer diagnosis, prognosis and therapy, and may pave pathways for the cure of solid and hematological cancers.

## Results

### Cytocapsular evolution analyses of breast cancers

To explore cytocapsular cancer evolution in breast cancers, we employed immunohistochemistry (IHC) staining and fluorescence microscope imaging analyses with anti-CM-01 antibodies that target the cytocapsular tube (CCT) marker protein CM-01, and with anti-gamma-actin antibodies and DAPI. We examined clinical normal breast tissues, benign breast tumor tissues, and 46 kinds of breast cancer tissues (3960 pieces of tissues) from the early stage, *in situ*, and stages I-IV from 3890 cancer patients. We analyzed >15,000 IHC fluorescence microscope images. We observed that in normal and benign breast tissues, cytocapsulae (CC) or cytocapsular tubes (CCT) are essentially absent, and CM-01 expression is very low (**Figs. S1A-B**). This indicates that normal and benign breast tissues do not generate CC/CCTs, and CM-01 expression is tightly controlled to a very low level in normal and benign breast tissues. In some highly compact early-stage breast tumors, we observed that there are neither CC nor CCT, CM-01 expression is very low, and intercellular space (ICS) percentage is 0% (**Fig. 1A**). These highly compacted transformed cells are tightly blocked by neighbor cells, preventing them to migrate, invade, or relocate (**Fig.1A**). These compact breast tumors are composed of transformed but incomplete cancer cells. They lose the normal biological function and structure of terminal ducts, and lack sufficient ICS (20%) for nutrient delivery and diffusion of metabolic waste molecules (**Fig. 1A**). These observations indicate that incomplete breast cancer cells are in a devolution process with multiple key features: lose normal breast cell segmentation capacity unable to form functional structures, unfastened normal breast cell biological functions, lacking normal cell contact inhibition, and gain of uncontrolled cell proliferation (**Figs. S1 and 1A**).

**Fig. 1.**
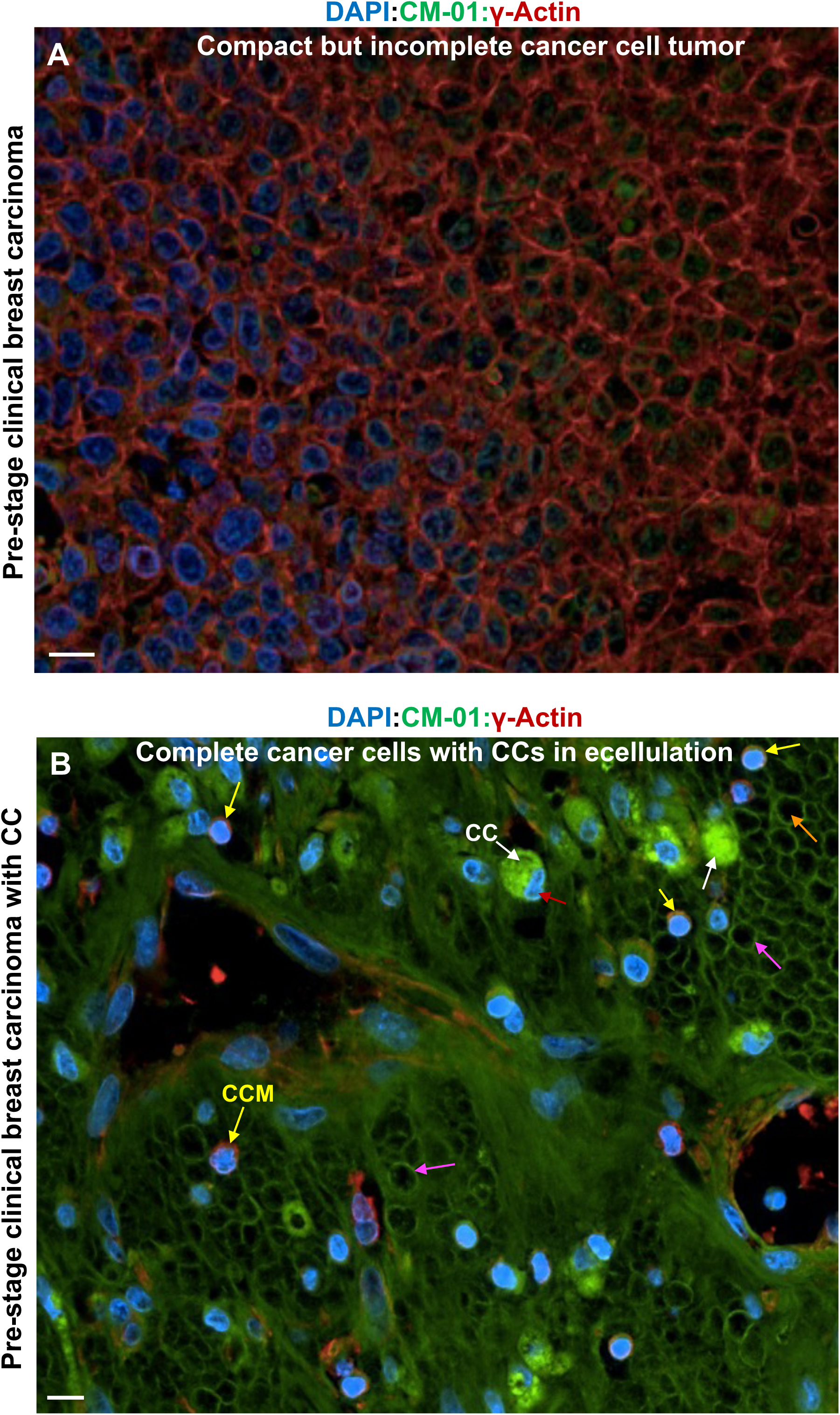
Transformed and incomplete cancer cells develop into complete cancer cells with cytocapsulae. (A) A representative immunohistochemistry (IHC) fluorescence microscope image of clinical breast carcinoma tissue composed of highly compact, transformed and incomplete breast cancer cells. The cytocapsular marker protein CM-01 is in low abundance. The transformed and incomplete breast cancer cells do not generate cytocapsulas (CC) or cytocapsular tubes (CCT). The intercellular space (ICS) is as low as 0% (normal tissue ICS is around 20%) due to the uncontrolled cell proliferation, and highly compact and dense tissue texture. The tightly compacted surrounding neighbor cells prevent transformed and incomplete cancer cells from migration, dissemination or escaping the local microenvironments. Scale bar, 10μm. (B) Representative IHC fluorescence microscope image of clinical breast carcinoma tissues in the very early cancer evolution stage. with complete cancer cells wrapped by cytocapsulae (CC, white arrows), complete cancer cells with cytocapsulas (CC) in the process of ecellulation, and ecellulated complete cancer cells. The complete cancer cells wrapped with small and tight CC membranes, complete cancer cells enclosed in bright green/gold color and folded CC membranes, complete cancer cells in the process of ecellulation, and many ecellulated CC are shown. CC membrane (yellow arrows) wrapping breast cancer cells, ecellulated CC (purple arrows) are shown. Scale bar, 10μm.

Interestingly, in some highly compact early-stage breast tumors, we observed that there are large quantities of CC/CCTs building blocks of tiny, membrane-enclosed vesicles in some transformed breast tumor cells’ plasma (**Fig. S1C**). There are no CCs or CCTs in the devolution stage of CC/CCT generation (**Fig. S1C**). Subsequently, these large quantities of tiny, membrane-enclosed vesicles are secreted outside the plasma membranes. They fuse together, and engender large CCs located outside the plasma membranes (**Fig. 1B**).

In some highly compact early-stage breast tumors, we observed that there are many cancer cell-generating folded membrane segments located outside the cells with high CM-01 abundance, indicating that these cancer cells engender CC outside the plasma membranes (**Fig.1B**). Surprisingly, there are large quantities of ecellulated CCs, and incomplete cancer cells left, leaving the ecellulated CC in compact and very dense textures (**Fig.1B**), which evidenced that ecellulation of CCs can massively occur in local areas in early-stage clinical cancer tissues *in vivo*. These observations are consistent with the *in-vitro* (in dishes) observations of CC ecellulation in thick 3D Matrigel. The above observations suggested that the generation of CCs at high abundance of CM-01 is tightly controlled, and serves for the physical separation of cancer cells from the stressful microenvironments and for the protection of cancer cells’ survival.

Next, we examined *in situ* breast cancer tissues of 3 subtypes from 12 patients, for which it was thought that breast cancer cells were present but not in a metastasis phase. Surprisingly, we found that there are large quantities of small CCTs in all of the checked *in situ* breast cancer tissues, and most of these breast cancer cell CCTs are very thin, of variable thickness (0.1-2μm in thickness), highly curved and of irregular morphologies but without degradation (**Fig.2A**). These observations suggest that these breast cancer cells generate CCs, breast cancer cells push CC membranes, elongate CC length, and engender thin CCT with a tail behind due to the strain from the moving cancer cell inside, and lack of anchoring of CCT in place in the ECM (extracellular matrix) by the nano-protrusion layers extension into the ECM (**Fig. 2A**). There are some CCTs already invading into the neighboring tissues. There are large quantities of breast cancer cell CCTs at the niches *in situ*. These observations indicate that cancer cell migration and invasion occur at the breast cancers *in situ* (**Fig. 2A**). Subsequently, breast cancer cells continuously push CCT membrane forward and elongate CCT length, and the nano-protrusion layers extend into ECM and anchor the CCTs in place. They shape the CCT morphologies together with the microfilament layer beneath the bi-layer lipid CCT membranes, and maintain the constant CCT diameter (**Fig.2B**). The neighboring transformed but incomplete breast cancer cells invade into these breast cancer cell CCTs, freely migrate in the membrane-enclosed physical freeways, and efficiently leave for far-distance destinations. Later, after most incomplete breast cancer cells have been moved by metastasis, these breast cancer cell CCTs degrade into strands of variable sizes, and followed by CCT strand decomposition and disappearance (**Fig.2B**).

**Fig. 2.**
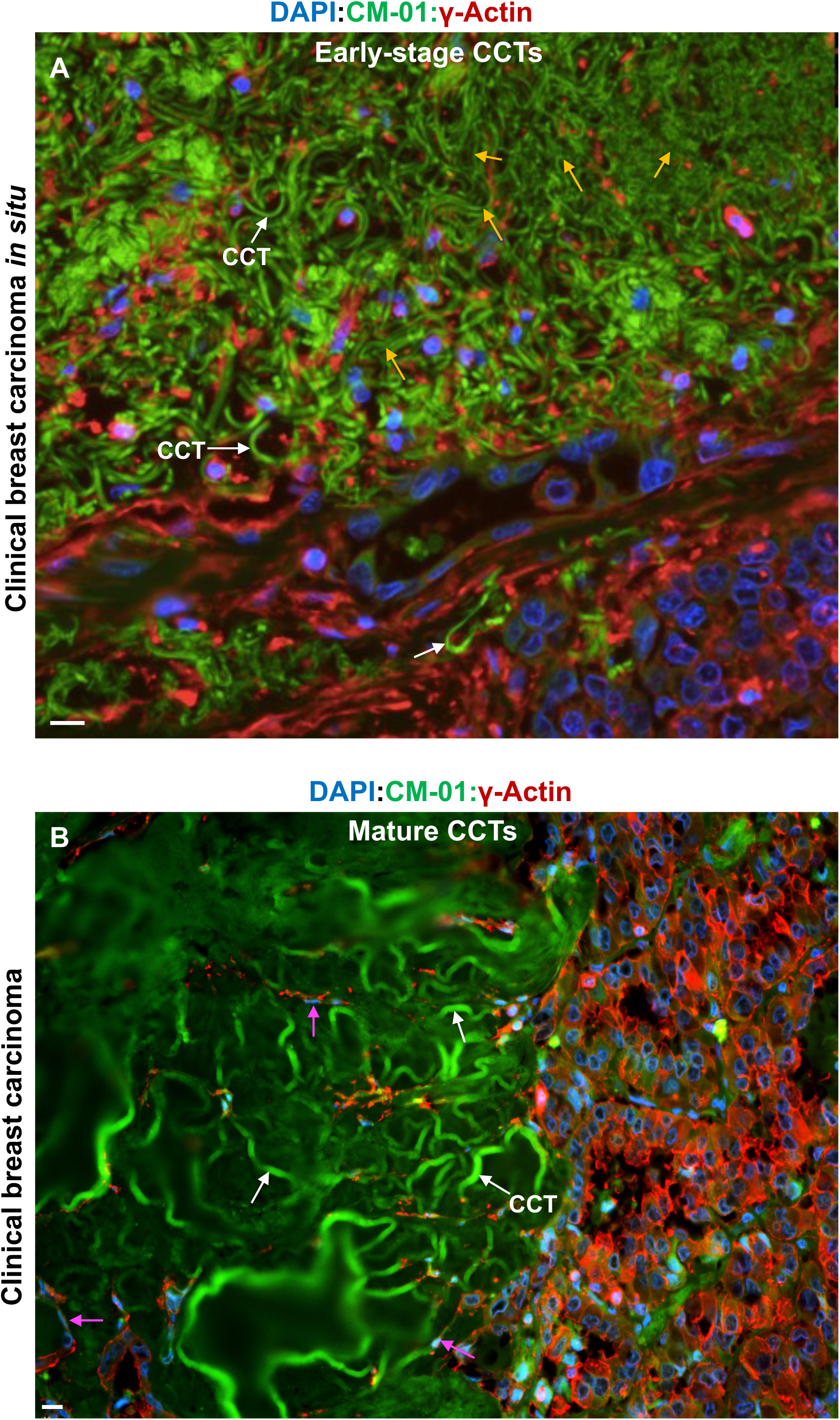
Cytocapsulae develop into initial thin and stretched CCT fragments, and then grow into mature long CCTs with constant CCT diameter. (A) Representative IHC fluorescence microscope image of clinical *in situ* breast carcinoma tissues. There are large quantities of cytocapsular tubes (CCT, white arrows) in the very early stage in the CCT lifecycle. Cancer cells in CC lumens secret large quantities of membrane enclosed vesicles, which fuse and integrate into CC membranes, and increase CC membrane sizes. Cancer cells push the folded CC membranes forward and leave a thin, long and stretched, tube-shaped vessels behind, which are CCTs, which are caused by the stretch of migrating cancer cells in CCT lumens, and lack of thick nano-protrusion layers surrounding the CCT fragment anchoring the CCT in place well. These thin and stretched CCT fragments (orange arrows) with inconstant diameters are not in degradation, and are different from the thin and long CCT strands which are caused by CCT membrane degradation. There are CCTs already invade into the neighbor tissues. Scale bar, 10μm. (B) Representative IHC fluorescence microscope image of mature CCTs in clinical breast cancer carcinoma. There are many long, highly curved, well anchored and mature and very long CCTs (up to 110 meters in length) in constant diameter (6um in measured diameter in this area), forming a CCT (white arrows) cluster and tightly contacting the compact tissues composed by many incomplete breast cancer cells. These incomplete breast cancer cells can invade into CCTs generated by other complete cancer cells, migrate inside, and leave for far away destinations. Breast cancer cells in migration in CCT (purple arrows) are shown. Scale bar, 10μm.

Interestingly, in the breast invasive ductal carcinoma tissues from 14 cancer patients, we found that many tumorspheres of various sizes and breast terminal duct-like strictures are wrapped in big or superlarge CC membranes. They are in various morphologies: round, oval, irregular shapes, with multiple circles or irregular shaped bubble-like morphologies. Most of these CCs are separated and isolated. Some CCs are fused/linked together to form complex structures (**Fig. S2A**). The invasive ductal breast cancer cells in the big CCs can continuously proliferate, differentiate, and generate terminal duct-like structures with abnormally thickened duct layers (2, 3, 4, or more layers of epithelial layer tissue), compared to single layers of epithelial cells in normal terminal ducts (**Fig. S2B**). These observations suggest that breast invasive ductal carcinoma cells can generate complete tumors in superlarge CCs, engender terminal duct-like structures with abnormal textures in superlarge CCs, and these CCs can separate from each other or fuse together to form complex structures (**Fig. S2**).

Subsequently, we found that the super large CC-wrapped compact breast malignant tumors grow into big sizes. The cancer cells in the complete compact malignant breast tumors can push the superlarge CC, extend the superlarge CC membranes in different directions and elongate the CCT lengths. Therefore, they generate many long CCTs surrounding the complete compact breast malignant tumors (**Fig. 3A**). Large quantities of CCTs, which originated from cancer cells inside the superlarge CC, surround and wrap the compact malignant tumors, and form thick CCT layers enveloping the malignant tumors in various thicknesses (from 3x to 30x folds of the compact malignant tumor sizes) (**Fig.3A**). All of these breast cancer-cell CCTs interconnect and generate 3D large CCT networks (**Fig.3B**). Breast cancer cells massively migrate in CCTs of these 3D large CCT networks (**Fig.3B**). In contrast, the non-invasive CT-Scan, MRI, or PET-CT imaging analyses, these tissues are classified as “normal” (actually are false normal tissues) due to the fact that cancer cell density in CCT networks is much less than the cell densities in compact tumors, and therefore are treated as “normal tissues” (**Fig.3B**).

**Fig. 3.**
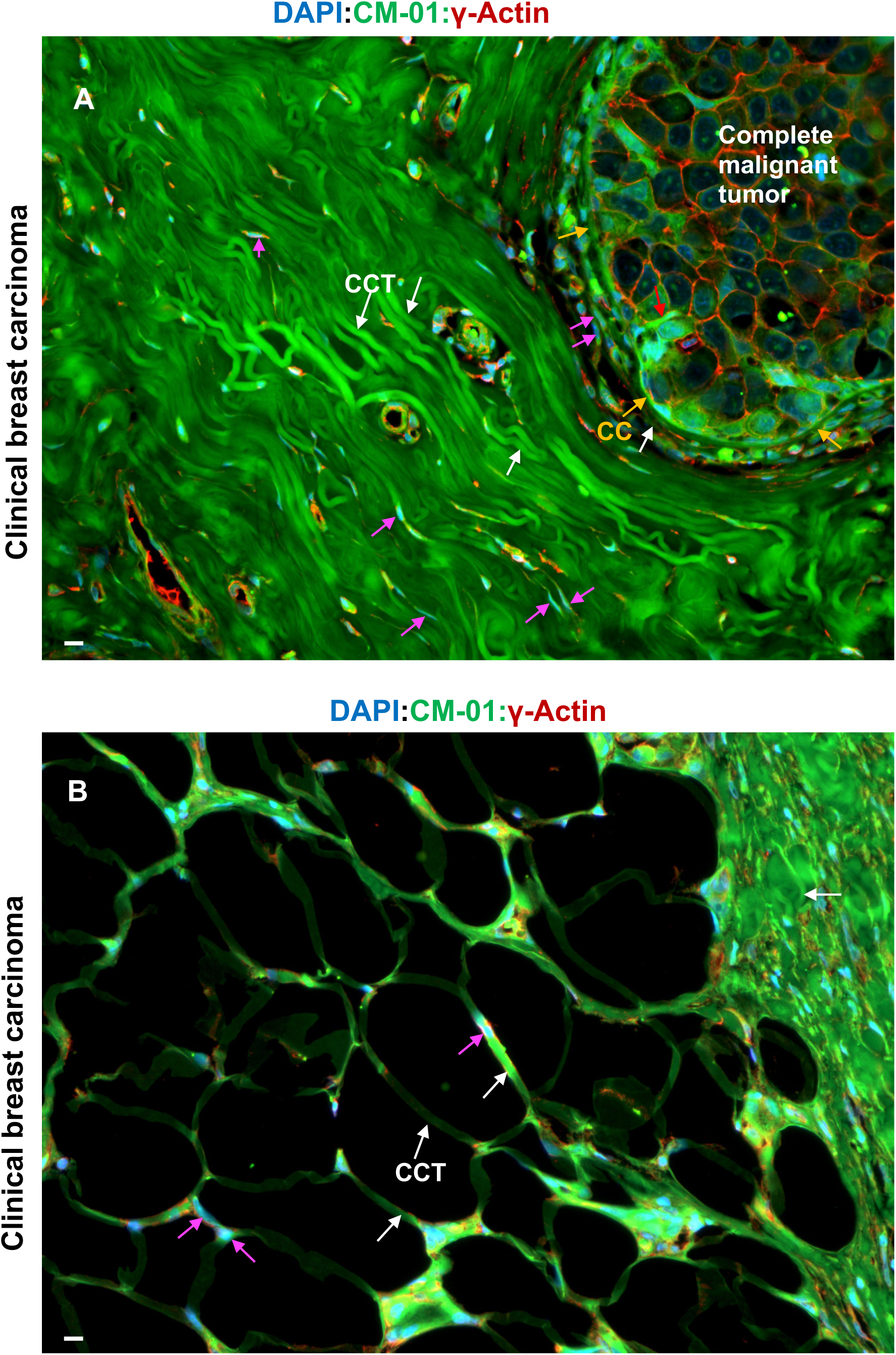
Development of complete malignant tumors and cytocapsular tube network-tumor system (CNTS). (A) Representative IHC fluorescence microscope image of a complete malignant tumor wrapped by thick CCT (white arrows) layers in clinical breast cancers. Single complete cancer cells divide and proliferate, and grow up into highly compacted and dense malignant tumor masses with 0% ICS; at the same time, cancer cells in the small CC secret large quantities of tiny membrane-enclosed vesicles, which fuse and integrate into CC membrane, and CC grow up into superlarge CC (orange arrows), wrapping the big compact tumor mass, and therefore generate complete malignant tumors. The cancer cells in the superlarge CC push, extend, and elongate superlarge CC membranes and generate many long CCTs, which wrap the compact complete malignant tumors many layers, and engender thick CCT layers enveloping the complete malignant tumors. Breast cancer cells in migration in CCT (purple arrows) are shown. A CCT (red arrow) generated by cancer cells in complete compact malignant tumor and is extending outside of the superlarge CC is shown. Scale bar, 10μm. (B) Representative IHC fluorescence microscope image of CCT networks in clinical breast carcinoma tissues. There are sectioned, large breast cancer CCT networks with breast cancer cells in migration in CCTs in the network. The right part is a compact, dense CCT network bunch which is composed of highly dense CCT networks. Scale bar, 10μm.

Compact malignant tumors in superlarge CC from the examined breast cancer patients are interconnected via large 3D CCT networks (**Figs. 3 and S3A**). The compact malignant tumors and the CCT networks originated from them, form a breast cancer CCT network-tumor system (CNTS) (**Figs. 3 and S3A**). Heterogeneous breast cancer cells freely migrate in the CNTS and can reach other heterogeneous malignant tumors in the CNTS (**Figs. 3 and S3A**), which increases the cellular heterogeneity of each compact malignant tumor in the CNTS. For the examined 1530 breast cancer patients with normal immunological functions, there are breast CNTSs composed of compact malignant tumors in CC and 3D CCT networks (**Figs. 3 and S3A, Table S1**). The coexistence of breast CNTSs and regular normal immune systems suggested that the CCT membrane enclosed CNTSs are recognized as normal self-tissues and escape the immune system attack. In the examined 1510 breast cancer autopsy tissue samples from 240 dead breast cancer patients that had been treated with multiple kinds of therapies (single kind therapy or combination therapies of surgery therapy, radiotherapy, chemotherapy, immunotherapy, gamma knife, proton knife), there are breast CNTSs composed of compact malignant tumors in CC and 3D CCT networks (**Figs. 3 and S3A, Table S1**). These observations suggested that breast CNTSs display potent drug resistance and damage resilience, and induce biological-functions failures to one or more essential organs, and therefore lead to breast cancer patient lethality (**Figs. 3 and S3A, Table S1**).

Consequently, small compact breast malignant tumors in superlarge CC grow up into medium and large sizes (**Fig. S3A**). The invasive ductal carcinoma cells generate terminal duct-like structures in superlarge CC continue to proliferate at a fast pace, and grow into compact malignant breast tumors wrapped in superlarge CCs (**Fig. S3B**). All of these compact malignant breast tumors of various sizes found in the patients are not separated or isolated from each other, but interconnect with each other via the 3D CCT networks composed of CCTs (**Fig. S3A-B, and Fig.3**). Breast cancer cells in the compact breast malignant tumors in the patient bodies can migrate in the 3D CCT networks and reach to other breast malignant tumors, or proliferate and grow into new breast malignant tumors in CCT terminals. Breast cancer cells in any site of the CCTs and 3D CCT networks can divide and grow into new breast malignant tumors (**Fig. S3A-B, and Fig.3**). They can go into apoptosis and necrosis (**Fig. S3B-C)**, and they can generate new CCTs and CCT networks within the compact complete breast malignant tumors. These CCTs can degrade, and CCT strands decompose, followed by CCT disappearance and intercellular fluid filling (**Fig. S3B-C**). These cavities in compact complete breast malignant tumors are the sources of tumor liquefaction (**Fig. S3B-C**).

Subsequently, after most cancer cells have invaded into CCTs and CCT networks, and left for far-distance destinations, CCTs will perform self-regulated CCT membrane degradation, and degrade into CCT strands of variable sizes (thick, thin, very thin), diverse morphologies (bunches, masses, cloud-like status) (**Fig. 4A**), and followed by CCT and CNTS decomposition and disappearance (**Fig. 4B**). These empty cavities caused by cancer cell metastasis and CCT decomposition will be filled by intercellular fluids, which are the resource of tissue liquefaction in tissues/organs in cancer patients at the late stages (**Fig.4B**). The cancer metastasis, disappearance of some compact malignant tumors in CC, decomposition of parts of CNTS, regeneration of new compact malignant tumors in CCs in the secondary niches of different tissues/organs, the broad interconnections among the compact malignant tumors developed in diverse tissues/organs and in different time, render the CNTSs in breast cancer patients spatiotemporally dynamic systems (**Figs 3-4, S2-3, and Table S1**). Therefore, breast CNTSs in breast cancer patients are spatiotemporal moving CNTSs, and its cytocapsular cancer evolution involves generation, growth, evolution, decomposition, and regeneration, in part or as a whole.

**Fig. 4.**
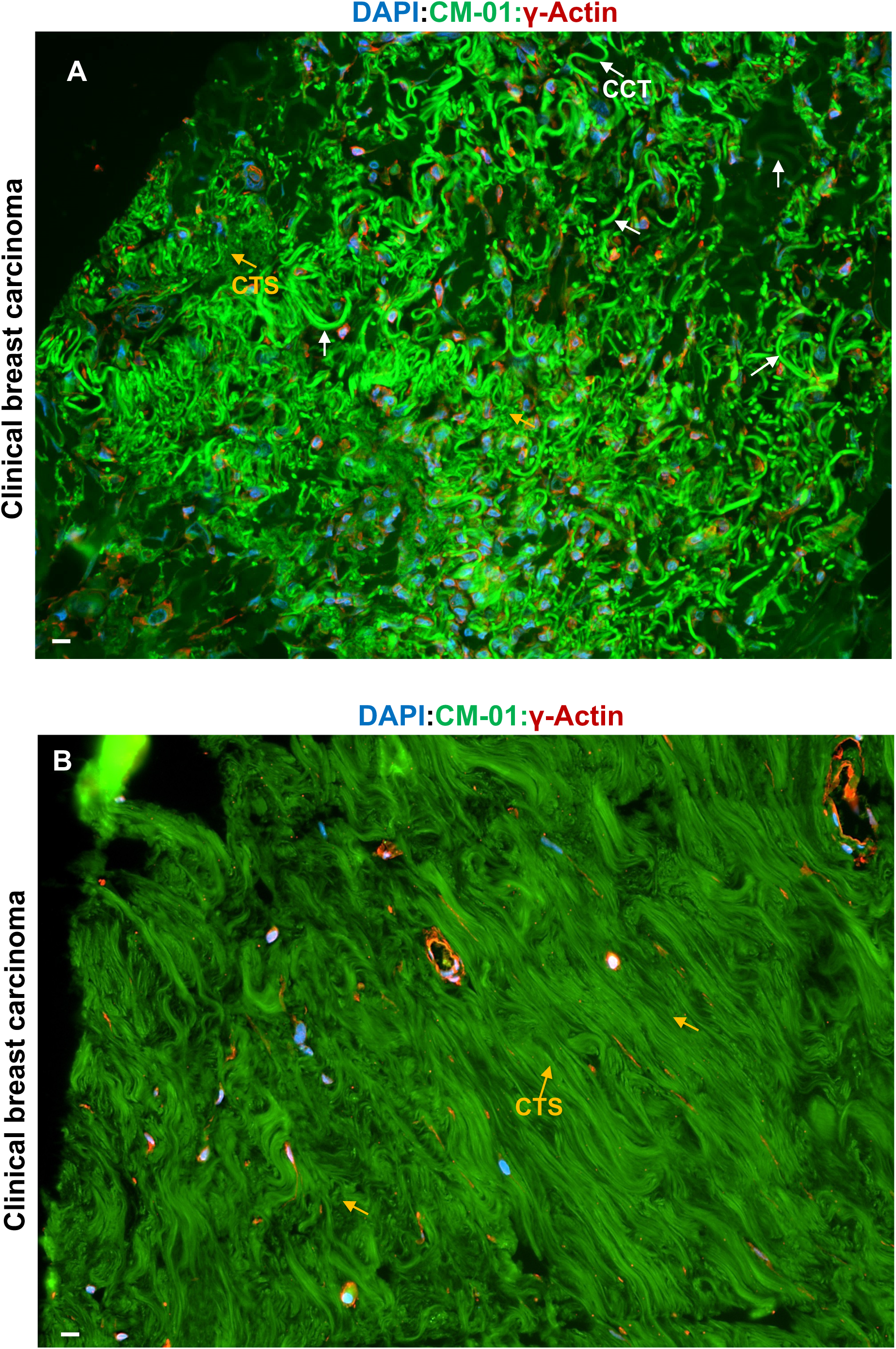
Cancer cell metastasis, degradation of CCTs, and decomposition of CCT networks and CNTSs. (A) Representative IHC fluorescence microscope image of degradation of CCTs (white arrows) in clinical breast carcinoma tissues. Cancer cells invade into CCTs and leave to far distance destinations. Most cancer cells (>98% in the imaged area) left and only a few remain, and cancer cell density significantly decreased. CCTs degrade into thick and thin strands (fragments). There is massive cancer cell CCT degradation and CCT network decomposed. Scale bar, 10μm. (B) Representative IHC fluorescence microscope image of very dense, severely degraded CCT strands (CTS, orange arrows), CCT networks, and CNTSs in clinical breast cancer tissues. Most (>98%) of cancer cells (within or beyond) compact malignant tumors have left to neighbor and far-distance tissues and organs, and there are only a few cancer cells remained. All the CCTs are degraded into very thin strands and form cloud-like CCT strand masses, compact malignant tumor cells disseminate, tumor metastasis, CCT networks decompose, and CNTS disappeared. Later, these CCT strands masses will continue to degrade into tiny debris, fragments, and completely disappear. Scale bar, 10μm.

A single piece of CC/CCT can be composed of many tiny membrane-enclosed vesicles secreted by multiple heterogeneous cancer cells, and shows heterogenous composition of the CC/CCT membranes. Heterogeneous CCTs can interconnect to form CCT networks, and CCT networks are propertied with heterogeneity in protein composition and abundance, lipid components and abundance and other abundant molecular components (**Figs. 1B, 2, 3, S2, and S3**). Heterogeneous cancer cells generate heterogeneous CC/CCTs, and CCTs in quite diverse CCT lifecycles. The data frequently show that, in the same tissue site, heterogeneous CCs, CCTs, and malignant tumors in CCs in different CC/CCT lifecycles coexist, displaying cytocapsular cancer evolution heterogeneity (**Figs.3, 4, S2 and S3**).

In summary, we discovered that breast cytocapsular cancer evolution in breast cancers includes: normal cell transformation, incomplete cancer cells, devolution and CC generation, complete cancer cells with cytocapsulae (CC), complete cancer cells in cytocapsular tubes (CCTs), complete malignant tumors in superlarge CC, CCT networks, cancer metastases, CCT network-tumor system (CNTS), CNTS decomposition and regeneration, and spatiotemporally dynamic CNTS. The heterogeneity broadly presents in breast cancer cell types/subtypes, CCs, sub-colony types/subtypes in CCs, breast malignant tumors in CC, CCTs, CCT networks, and breast CCT network-tumor systems (CNTS).

### Cytocapsular evolution analyses of brain cancers

To explore cytocapsular cancer evolution in brain cancers, we investigated normal brain tissue (cerebellum from monkey), and 3 subtypes of brain cancer tissues from 122 brain cancer patients in stages I-IV, including one of the most aggressive and deadly brain cancers of glioblastoma (GBM). We observed that normal neurons cells in cerebellum have many long dendrites and the CC/CCT marker protein CM-01 is at a very low level (**Fig. S4A**). In contrast, we found that transformed and incomplete GBM cancer cells do not have long/short dendrites or synapses, and exhibit a highly compacted texture with the intercellular space (ICS) percentage as low as 0% (**Fig. S4B**). The CCT marker protein CM-01 in some incomplete GBM cancer cell fragments are at high abundance, but displaying CM-01 abundance heterogeneity. The incomplete GBM cancer cells at this stage do not generate CCs outside of the GBM cell membranes (**Fig. S4B**). These observations indicate that incomplete GBM cancer cells go into a devolution process with multiple key phenomena: lose normal neuron cells’ structures of dendrites and synapses, lack normal neuron functions, lose cell contact inhibition, and gain uncontrolled cell proliferation (**Fig. S4**).

Interestingly, in the early stage GBM tissues, in parts of the tissues, a lot of incomplete GBM cancer cells generate cytocapsulas (CC) outside of the GBM cancer cell cytoplasm membranes in various sizes (small, medium and large) (**Fig. S5A**). The GBM cancer cells with CC are complete GBM cancer cells. The CC sizes of GBM cancer cells are statistically larger than those of breast cancer CC (**Fig. 1B and Fig. S5A**). These complete GBM cancer cells with CC are usually in dense tissue texture (**Fig. S5A**).

Subsequently, GBM cancer cell push CC, and elongate CC membranes and generate long GBM cancer cell CCTs (**Fig. S5B**). The long GBM cancer cell CCTs can potently invade into neighboring or far-distance highly compact and dense GBM tumor tissues (**Fig. S5B**). GBM cancer cells invade into and migrate in GBM cancer cell CCTs, leave for far-distance destinations, and lead to decreased cell density in local niches. Many GBM cancer cell CCTs interconnect and form 3D CCT networks (**Fig. S6A**).

Consequently, after most of GBM cancer cells left, GBM cancer cell CCTs and CCT networks degrade into variable strands (thick, thin, very thin) in quite diverse morphologies (curved, coiled, winded, irregular shaped) (**Fig. S6B**). The cell density in the local niche significantly decreases and only few GBM cancer cells remain (**Fig. S6B**). Later, GBM cancer cell CCT strands continue to degrade into very thin strands in large quantities in a cloud-like status (**Fig. S7A**). Due to the massive GBM cancer cell CCT invading into micro blood vessels, micro blood vessels are broken and leaky, and many red blood cells are released and randomly distributed into the local tissues (**Fig. S7A**). Subsequently, very thin strands of GBM cancer cell CCTs decompose and disappear. More micro blood vessels are broken and damaged, and red blood cells are released and randomly distributed in the local tissues (**Fig. S7B**). Therefore, after GBM cancer cells metastasized and left, GBM cancer cell CCTs degrade and decompose, local micro blood vessels are broken, and the brain tissue in these areas lose normal biological functions, which leads to failure of brain local tissue biological functions (**Fig. S7**).

In summary, we discovered that cytocapsular cancer evolution in brain cancers share most of the features of that in breast cancers (**Figs 1-4, and S1-7**).

### Cytocapsular evolution analyses of other 263 kinds/subtypes of solid cancers

To comprehensively understand cytocapsular cancer evolution in other 263 kinds/subtypes of solid cancers (**Table S1**), we investigated 13 subtypes of pancreas cancer tissues from 218 pancreas cancer patients in all stages I-IV (**Fig. S8**), 5 subtypes of thyroid cancer tissues from 17 thyroid cancer patients in all stages I-IV (**Fig. S9**), 5 subtypes of oral cancer tissues from 17 oral cancer patients in all stages I-IV (**Fig. S10**), 25 subtypes of prostate cancer tissues from 1528 prostate cancer patients in all stages I-IV (**Fig. S11**), 23 subtypes of lung cancer tissues from 676 lung cancer patients in all stages I-IV (**Fig. S12**), 12 subtypes of bone cancer tissues from 210 bone cancer patients in all stages I-IV (**Fig. S13**), 2 subtypes of smooth muscle cancer tissues from 20 smooth muscle cancer patients in all stages I-IV (**Fig. S14**), 2 subtypes of fibrous cancer tissues from 25 fibrous cancer patients in all stages I-IV (**Fig. S15**). In short, we found that cytocapsular cancer evolution in 265 kinds/subtypes of solid cancers (**Figs. 1-4, S1-15, Table S1**) includes multiple successive devolution and evolution procedures: 1) devolution includes normal/neoplasia/nodule cell transformation, and CC/CCT generation; 2) evolution consists of: complete cancer cells in CCTs, complete malignant tumors in superlarge CC, CCT network formation, cancer metastasis, secondary complete malignant tumor formation and growth, CCT network-tumor system (CNTS), CCT degradation and decomposition, CNTS regeneration, moving CNTS, and moving damages to normal tissues (**Fig.5**). The heterogeneity broadly displays in solid cancer cell types/subtypes, CC, sub-colony types/subtypes in CC, malignant tumors in CC, CCTs, CCT networks, and CNTSs (**Fig.5**).

**Fig. 5.**
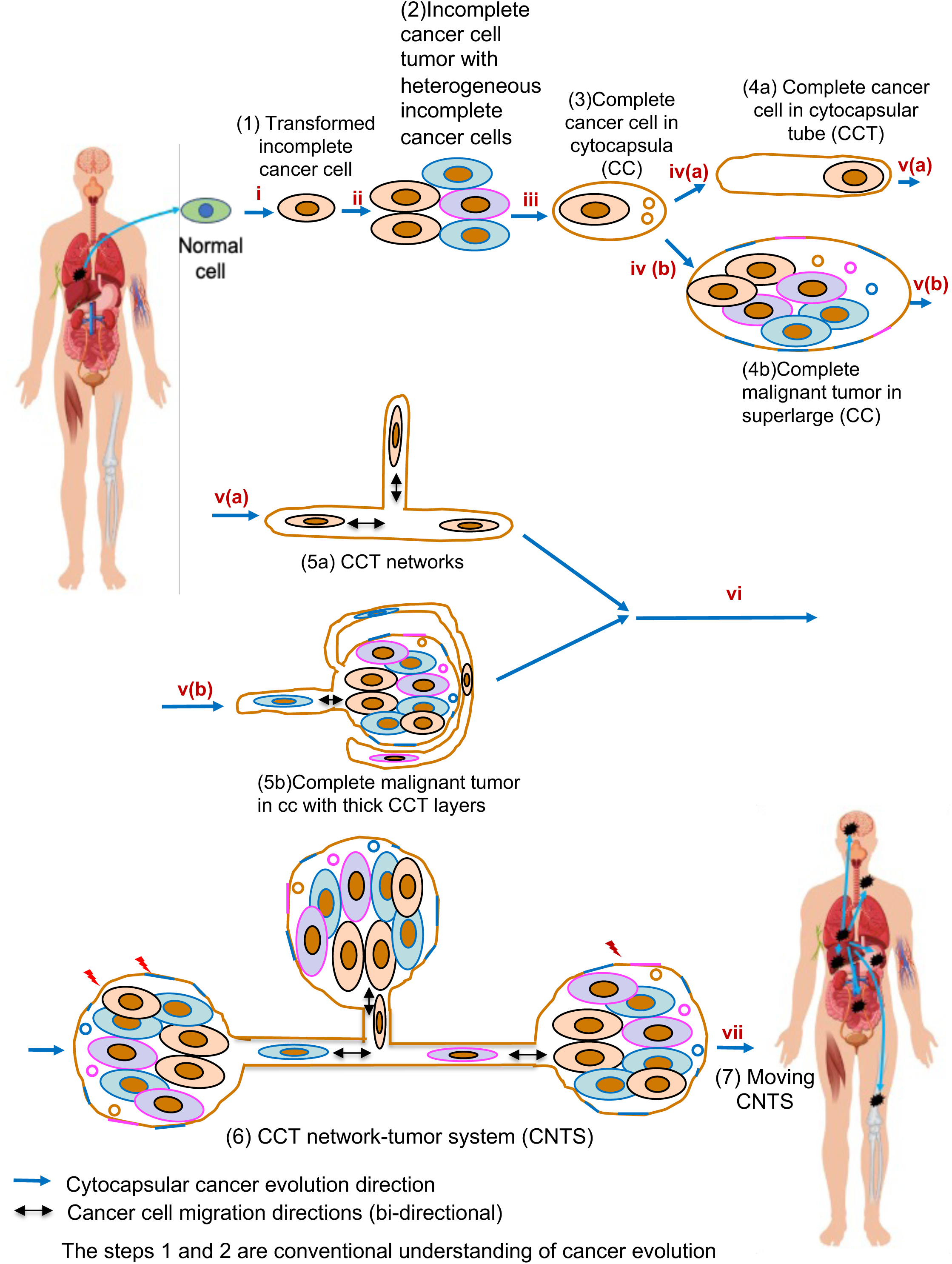
A schematic diagram of cytocapsular cancer evolution pathway of solid cancers. There are 7 major successive phases in the cytocapsular cancer evolution pathway of solid cancers: i) Devolution and transformation. Normal cell (or neoplasia cell, nodule cell) is devolved and transformed into transformed but incomplete cancer cell (1) with accumulated gene mutations and stressful ECM stimuli. The devolution procedures display some key features: lose normal breast cell segmentation capacity unable to form functional structures, unfastened normal breast cell biological functions, lacking normal cell contact inhibition, and gain of uncontrolled cell proliferation. ii) Uncontrolled proliferation. Transformed but incomplete cancer cells undergo uncontrolled cell proliferation, and generate incomplete cancer cell tumor/mass with heterogeneous incomplete cancer cells (2), which lead to nutrient deprivation, hypoxia, low *p*H, elevated toxic metabolic molecules, and stressful microenvironments. These incomplete cancer cells have to face two options: apoptosis or try to survive. iii) Devolution and cytocapsula (CC) formation. Incomplete cancer cells experience devolution and generate cytocapsulas (CCs), and engender complete cancer cells. iva) Complete cancer cells secret tiny vesicles, which integrate into CC and increase CC membrane size. Cancer cells in CC push CC, and elongate CC, and form cytocapsular tube (CCT) (4a). va) CCT perform branching morphogenesis and generate CCT networks. Cancer cells freely migrate in CCT networks (5a). ivb) Complete cancer cells divide, proliferate, and generate compact tumor in the superlarge CC, and form compact complete malignant tumor (4b). vb) Cancer cells in complete malignant tumors can push the superlarge CC membrane, and generate CCTs extending outside, or wrapping the complete malignant tumors. Cancer cells in complete malignant tumors can also generate its CCTs, and extend outside, and interconnect other CCTs and form CCT networks outside the complete malignant tumors. The CCT networks wrap the complete malignant tumors in various thickness, and form CCT network-tumor complex (5b). vi All the compact complete tumors in cancer patient bodies are interconnected via CCT networks, and form a CCT network-tumor system (6). All complete heterogeneous malignant tumors in cancer patients are not separated or isolated, but interconnect to each other via 3D heterogeneous CCT networks composed of heterogeneous CCTs. Heterogeneous cancer cells freely and efficiently migrate in membrane-enclosed CCT network freeway systems crossing tissues and organs. The evolution of cancer cells, CCTs, CCT networks, complete malignant tumors integrate together and drive the CCT network-tumor system (CNTS) evolution. Immune cells recognize CCT membranes as self-tissues and cancer cells and tumors in CC/CCTs escape immune therapy, displaying immune-therapeutical cold tumors. CCT membrane barrier leads to pan-resistance to conventional cancer drugs. vii) Moving CNTS: CNTSs are spatiotemporally dynamic. CNTS generation, decomposition, and regeneration, and metastasis to diverse tissues and organs in different time. CNTS displays dynamic features in: CCT generation, degradation, decomposition, and new CCT generation; complete tumor generation, apoptosis, growth, metastasis, and new complete tumor growth; CCT network generation, decomposition, and regeneration; cancer cell metastasis; new CNTS in new normal tissues and organs. The moving CNTSs brining moving damages and biological function failure to normal tissues and organs.

### Cytocapsular evolution analyses of 46 kinds of hematological cancers

To explore cytocapsular cancer evolution in liquid (hematological) cancers, we investigated 46 kinds/subtypes of liquid (hematological) cancers (**Fig. S16 and Table S1**). We found that cytocapsular cancer evolution in liquid cancers contains: cancer transformation, incomplete cancer cells, devolution and CC generation, complete cancer cells with cytocapsulas (CC), complete cancer cells in cytocapsular tubes (CCTs) and CCT network in bone marrow, thymus, lymph nodes, and spleen, but only with CC in the blood, CC degradation and decomposition, regeneration of CC, CCTs and CCT networks in bone marrow, thymus, lymph nodes, and spleen, moving CCT networks, moving damages to normal tissues in bone marrow, thymus, lymph nodes, and spleen (**Fig. S16 and Table S1**).

### An atlas of cytocapsular cancer evolution

Combined all of the above analysis data together, we created an atlas of cytocapsular cancer evolution of solid and liquid (hematological) cancers (**Fig. 6**). There are 6 major steps in the cytocapsular cancer evolution in solid cancers in human *in vivo*:

i) Transformation: transformation of normal cells/neoplasia cells/nodule cells/benign tumor cells into incomplete cancer cells (no CC) in normal extracellular matrix (ECM) leads to the latter in a devolution process: lose normal cells’ structures, functions and capacities, lose cell contact inhibition, gain of uncontrolled cell proliferation;
ii) Uncontrolled cell proliferation: uncontrolled cell proliferation of incomplete cancer leads to highly dense tissue textures, decreased intercellular space (ICS), loss of nutrient delivery pathways, nutrient deprivation, insufficient growth factors, enhanced metabolic wastes, elevated toxic molecules, local hypoxia, low *p*H, and stressful microenvironments;
iii) Devolution and CC generation: the stressful microenvironments stimulate the incomplete cancer cells to activate the ancient capacity and secrete a lot of membrane-enclosed tiny vesicles outside of and close to the cell membrane; these tiny vesicles fuse together and form a large bi-layer lipid membrane, biomembrane-enclosed cytocapsulae outside the incomplete cancer cells, and generate CC/CCT outside of the cell membrane, wrap the incomplete cancer cells inside, physically separate incomplete cancer cells from the stressful microenvironments; this devolution procedure creates a secondary membrane vessel outside of the cell membrane and protects the incomplete cancer cells for survival, and engender a complete cancer cell with both key characters: uncontrolled cell proliferation, and cancer cell metastasis via physical CCT freeways for escaping towards better microenvironments for continuous cell survival; individual cancer cells in CC or CCT can go into a cellular dormancy status;
iv) Cancer cell proliferates in CC/CCTs: form malignant tumors in superlarge CC; the complete malignant tumors can collectively go into a tumor mass dormancy status;
v) CCT branching morphogenesis: CCT network formation; CCT networks interconnect all the primary and secondary tumors in metastatic sites (including bone marrow and brain);
vi) CNTS formation: Interconnection of all tumors via CCT networks generates CCT network-tumor systems (CNTSs); with conventional cancer drug pan-resistance and damage resilience properties across tissues and organs in cancer patient bodies
vii) Spatiotemporally dynamic CNTS: CNTS decomposition and regeneration in primary and secondary niches present moving CNTS or spatiotemporally dynamic CNTS, and induce moving damages in multiple tissues/organs.

**Fig. 6.**
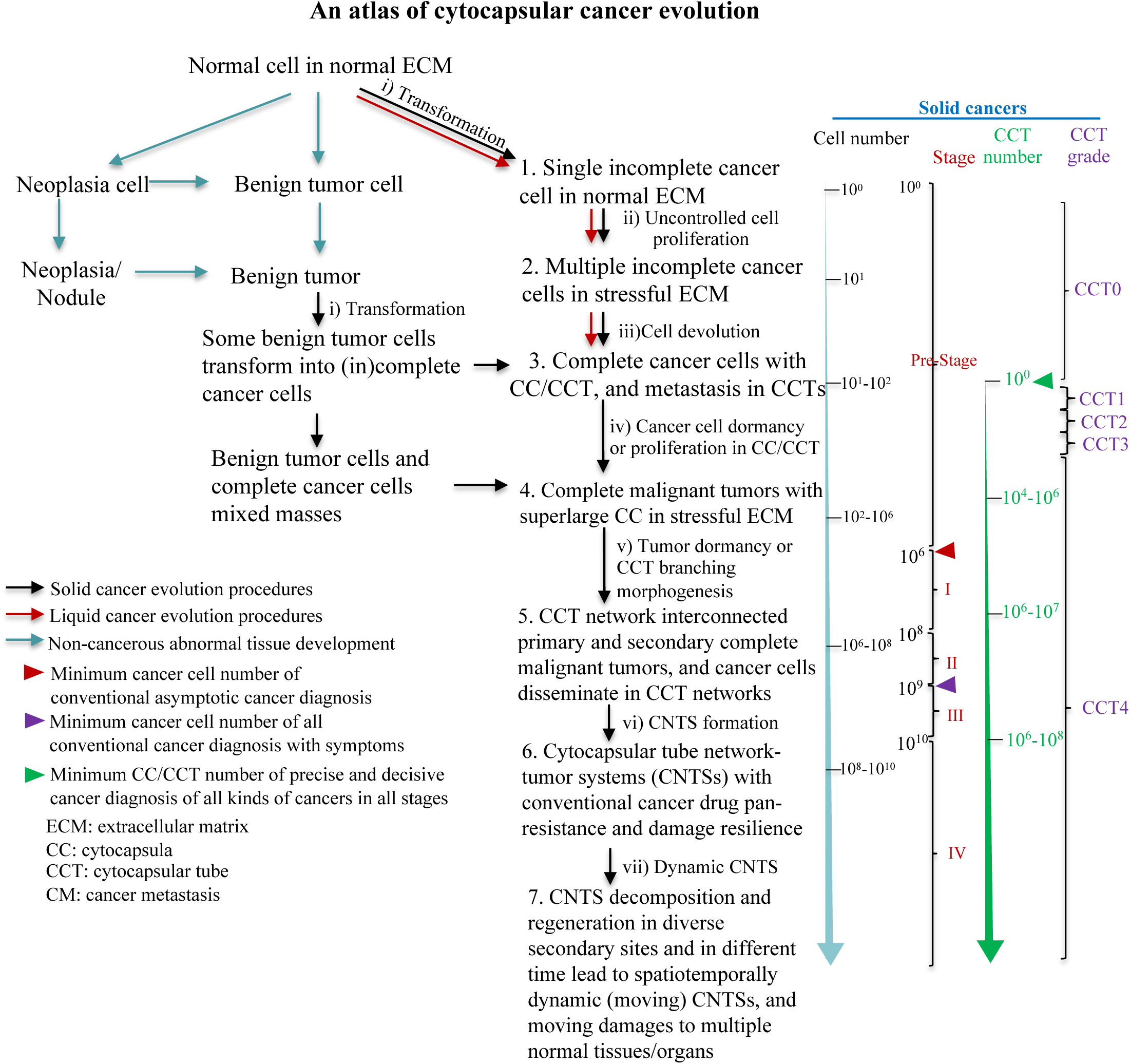
A cytocapsular cancer evolution atlas. Cytocapsular cancer evolution pathways in solid cancers are shown in black arrows, and that in liquid (hematological) cancers are presented in red arrows. The non-cancerous evolution activities are shown in green arrows. In the left part, the long and cyan color arrow shows the rough cancer cell number of each phase of the 7 phases. The words in red show the conventional cancer stages (pre-stage, and stages I, II, III and IV) and the black braces show the rough cancer cell numbers of each cancer stage. The long and green arrow shows the roughly estimated CCT numbers in the cytocapsular cancer evolution phases 3-6 in cancer patients. The words in purple show the CCT grades (CCT0, CCT1, CCT2, CCT3, and CCT4) with the purple braces showing the CCT number ranges. The details of CCT grades are shown in a previous article^15^.

For liquid (hematological) cancers, there are 3 major steps in cytocapsular cancer evolution in human *in vivo* (red arrows in **Fig. 6**):

i) Transformation: transformation of normal blood and lymph cells into incomplete cancer cell (no CC) in normal microenvironments;
ii) Uncontrolled cell proliferation: uncontrolled incomplete cancer cell proliferation leads to stressful microenvironments in bone marrow/lymph node/spleen/thymus (or occasionally in blood with elevated stressors, and increased viscosity caused by elevated lipids, fats, proteins, or cell fragments, or combination);
iii) Devolution and CC generation: incomplete liquid cancer cells generate CC/CCTs, and engender complete liquid cancer cells;
iv) Cancer cell proliferation in CC, CCTs: complete liquid cancer cells proliferate in CC, CCTs, and CCT networks in bone marrow/ lymph node/spleen/thymus (no CCT or CCT network in the circulation), and then, liquid cancer cells are released into the circulation via CCT invasion into micro blood vessels (CC may degrade in normal microenvironments or remain/regenerate in stressful microenvironments). Liquid cancer cells homing in bone marrow/lymph node/spleen/thymus will regenerate CCT and CCT networks.

## Discussion

The understanding of the nature of cancer evolution is a prerequisite and essential for all aspects of cancer prevention, control, therapy and cure^20–25^. This study investigated cytocapsular cancer evolution of 311 kinds /subtypes of clinical cancers, and elucidated an atlas of cytocapsular cancer evolution *in vivo* of both solid and liquid (hematological) cancers.

Based on our study of cytocapsular cancer evolution, together with recent advancements in molecular evolution, and cancer evolution hypotheses, we therefore propose 5 key features of cytocapsular cancer evolution: 1) Heterogeneity; 2) Heredity; 3) Devolution and evolution; 4) Selection and adaption/death; and 5) Metastasis and moving survival/damage. Heterogeneity presents in gene mutation, genomic aberrations, metabolic addiction, incomplete and complete cancer cells, cell plasticity, complete tumors in CC, CCTs, CCT networks, and CCT network-tumor system (CNTS). Heredity presents in genome, biological information flow principles, cancer cell types, loss and acquired biological functions of cancer cells. Devolution and evolution: devolution includes: transformation of normal/neoplasia/nodule cells, and generation of CC/CCTs; evolution consists of: grow into complete malignant tumors, CCT network formation, CNTS generation, moving CNTSs. Selection and adaption/death contains: stressful microenvironment selection, interference/therapy selection, adaption to survival, adaption failure leading to cancer cell death. Metastasis and moving survival/damage consist of: cancer metastasis for continuous survival (moving survival) at both individual and sub-colony/colony levels, and bring moving damages (biological functional failure) to multiple normal tissues/organs.

The presence of CC, CCTs, CCT networks, and CNTS significantly enhances cancer cell capacities in multiple aspects: 1) devolution and evolution; 2) selection and adaption; and 3) metastasis and continuous (moving) survival. At the same time, it significantly increases the difficult for cancer therapy at multiple levels: 1) CC/CCT membrane drug barrier, 2) CNTS complex system, 3) spatiotemporal moving CNTS (CNTS generation, decomposition, and regeneration), 4) heterogeneity in CCT composition components, density, lifecycle and super-structures; 5) inherent and acquired therapy/interference resistance; 6) inherent and acquired resilience to CNTS physical damages (the remained CNTSs still function and relapse); 7) bring the moving damages to multiple normal tissues/organs.

The discovery of cytocapsular cancer evolution pathways and atlas expands our understanding of the nature of cancer devolution and evolution. The discoveries described in this study extend our understanding with: complete cancer cells with CC/CCT, complete malignant tumor in superlarge CC and with thick CCT layers, CCT networks, CNTS, and the heterogeneity in complete cancer cells, complete malignant tumor in superlarge CC, CCT networks, CNTS, spatiotemporal moving CNSTs, moving damages. The identification of CCT addicted protein marker CM-01 in 311 kinds/subtypes of cancers suggests that it is possible to target CCT marker genes/proteins for the elimination and eradication of both CCTs and cancer cells in CCTs. The molecular mechanisms of each step of cytocapsular cancer evolution are largely unknown, and need more research to elucidate.

In summary, this study discovered cytocapsular cancer evolution pathways and established an atlas for both solid cancers and liquid (hematological) cancers, which will facilitate cancer research, cancer diagnosis, cancer drug discovery and development, and cancer therapy. This in turn may pave avenues towards the cure of all kinds of cancers.

## Methods

### Materials and Reagents

The patients providing formalin-fixed paraffin-embedded tissue samples gave informed consent that they understood that the biopsies (needle biopsy or surgical biopsy, from US Biomax) were performed for *in vitro* research purposes only. Comparative deidentified samples of normal tissues, benign tissues, carcinoma *in situ*, cancer, paracancer, metastatic tissues with their cancer stages identified according to the tumor (T), node (N), and metastasis (M) TNM system (cancer stages: 0, I, II, III, IV) were obtained from archival materials. The cancer, paracancer and metastatic cancer tissues, in which the original cancer niches were identified by indicated cancer specific molecular markers, were identified by hospital pathology laboratories and obtained from archival material.

### Histology and Immunohistochemical staining analysis

Cancer lesion samples were obtained from the index patients from clinical hospitals in USA (356 specimens from US Biomax; others samples directly from clinical hospitals). Samples were processed for histopathological evaluation with the use of hematoxylin and eosin staining. Immunohistochemical fluorescence tests were performed to stain cytocapsular tubes using rabbit anti-CM-01 monoclonal and polyclonal primary antibodies (self-made; CM-01: a code, not real protein name), mouse anti-gamma-actin monoclonal primary antibodies (Abcam, ab123034), Goat anti-Mouse IgG (H+L) Highly Cross-Adsorbed Secondary Antibody, Alexa Fluor Plus 555 (Thermo Fisher), and Goat anti-Rabbit IgG (H+L) Highly Cross-Adsorbed Secondary Antibody, Alexa Fluor Plus 488, Thermo Fisher).

### Data collection

Cytocapsular tubes (CTs, not sectioned, longitudinally sectioned, and cross sectioned) without degradation (3∼6μm in measured diameter) were counted using a fluorescence microscope and ImageJ. The presence of CTs degrading into thick strands (1∼2μm in measured diameter), thin strands (0.2∼1μm in measured diameter), or the disintegration state were reported without quantification.

### Statistical analysis

For each specimen, the number of fully intact cytocapsular tubes was counted in 5 areas (0.35mm x 0.35mm, length x width) of the sample (top, bottom, left, right, and center), and the cytocapsular tube density (CT/mm^2^) was calculated and determined for each area. The average CT density across the 5 sites was treated as the specimen’s overall CT density.

Supported by a grant (to Dr. Yi) from Cytocapsula Research Institute Fund for Cytocapsular Tube Conducted Cancer Metastasis Research, and a grant (to Dr. Yi) from Cellmig Biolabs Inc Fund for Cytocapsular Tube Cancer Metastasis Research. All data produced in the present work are contained in the manuscript.

We greatly acknowledge Dr. Ed Harlow of Harvard Medical School (USA) and Cancer Institute of University of Cambridge (UK), and Dr. Nahum Sonenberg of McGill University of Canada, Dr. Yubo Yang and Dr. Qiping Hou for their help and meaningful discussion in the study.

## Supplementary information

**Fig. S1. Comparison of tissue textures of clinical normal breast tissue, clinical breast benign tumor, and clinical breast carcinoma in early stage.**

(A) Representative fluorescence microscope image of normal breast tissue with immunohistochemistry (IHC) staining with anti-CM-01 (green), anti-gamma actin (red) antibodies and DAPI. The expression of CM-01 in normal breast tissue is in a low abundance. There are multiple terminal ducts (TD) with single epithelial cell layer. The statistical average intercellular space (ICS) percentage is about 20% (n=5 pieces of different normal breast tissues). Scale bar, 10μm.

(B) Representative fluorescence microscope image of clinical breast benign tumor tissues. The expression of CM-01 in breast benign tumor tissue is in a low abundance. There is no terminal duct. The breast benign tumor tissue is composed of abnormal cell masses with intercellular space (ICS) variably from 4%-6%. Scale bar, 10μm.

(C) Representative fluorescence microscope image of clinical breast carcinoma in the early stage. There is no terminal duct. The highly compacted and dense tissue texture with ICS percentage as low as 0%. In the cellular plasma of some breast cancer cells, there are many tiny, membrane-enclosed vesicles with high abundance of CC/CCT marker protein CM-01 (white arrows), which are building blocks of CC/CCT membranes. There is no cytocapsula (CC) generated yet. Scale bar, 10μm.

**Fig. S2. Complete cancer cells in cytocapsulae (CC) proliferate and evolve into malignant tumors and cancerous complexes in diverse morphologies.**

(A) Representative fluorescence microscope image of clinical breast invasive ductal carcinoma. Single complete cancer cells in CC divide, proliferate, differentiate, and evolve into compact tumorspheres in small, big and all kinds of variable sizes, and terminal duct-like structures with abnormal thickness and sizes, in which all tumorspheres and abnormal duct-like structures are enclosed in the superlarge cytocapsulae (CC, orange arrows) in quite diverse shapes and morphologies. Many diverse carcinoma texture structures in superlarge CC are shown. Scale bar, 10μm.

(B) Single complete cancer cells in CC proliferate, differentiate and evolve into big and complex, and abnormal duct-like structures, which are wrapped in superlarge CC (orange arrows). Several abnormal duct-like structures enveloped in superlarge CC are shown. Scale bar, 10μm.

**Fig. S3. Complete malignant tumors, massive apoptosis/necrosis in complete malignant tumors, and CCT network degradation in complete malignant tumors in clinical breast carcinoma.**

(A) Representative fluorescence microscope image of multiple small and middle-sized malignant tumors wrapped in superlarge CC (orange arrows) in cytocapsular tube network-tumor system (CNTS). There are many CCTs and CCT networks surrounding these complete malignant tumors, which are dimmed due to unfocused layers and/or CCT degradation. Scale bar, 10μm.

(B) Representative fluorescence microscope image of massive apoptosis/necrosis in complete compact malignant tumors enclosed in superlarge CC (orange arrows). In the complete compact malignant tumor at the center of the panel, there are many cancer cells in apoptosis/necrosis. This complete compact malignant tumor is a tumor part of a CNTS. Scale bar, 10μm.

(C) Representative fluorescence microscope image of CCT networks, CCT network degradation within the complete compact malignant tumors which are wrapped in superlarge CC (orange arrows), and parts of CNTS in clinical breast carcinoma. In the complete compact malignant tumors, some cancer cells generate CCTs and CCTs (white arrows); these CCTs extend outside of the superlarge CC, and expand into neighbor and far-distance destinations. There are multiple complete compact malignant tumors with diverse textures inside: 1) compact tumor masses, 2) CCTs and CCT networks that facilitate cancer cells inside the complete compact malignant tumors to leave away and reach far destinations beyond these complete compact malignant tumors, 3) massive cancer cell apoptosis occurrence within complete compact malignant tumors. The phenomena in the above items 2) and 3) are “tumor liquefaction” in malignant tumors. Scale bar, 10μm.

**Fig. S4. Increased abundance of CC market protein CM-01 in some incomplete glioblastoma (GBM) cells in cerebellum in clinical early stage GBM tissues.**

(A) Representative fluorescence image of monkey normal cerebellum tissue (as a control). All normal neurons cells in cerebellum have many dendrites and the CC market protein CM-01is in low abundance. Scale bar, 10μm.

(B) Representative fluorescence image of human clinical GBM in the early stage. All transformed and incomplete GBM cancer cells do not have long dendrites or synapses, and they are in highly compacted texture with ICS percentage as low as 0%. The CC/CCT marker protein CM-01 in some incomplete GBM cancer cells are in high abundance, but these incomplete GBM cancer cells do not generate CC outside of the GBM cell membranes yet in this imaged field. Scale bar, 10μm.

**Fig. S5. Complete glioblastoma (GBM) cancer cells generate cytocapsulae (CC) and cytocapsular tubes (CCTs) in the clinical early stage GBM tumors.**

(A) Representative fluorescence image of complete GBM cancer cells with CC. All complete GBM cancer cells in this area generate small, middle or big CC (white arrows) located outside of GBM cell membranes. All these complete GBM cancer cells do not have long dendrites or synapses. These complete GBM cancer cells are in compact and dense texture and the ISC percentage in this imaged area is as low as 0%. Scale bar, 10μm.

(B) Representative fluorescence image of GBM cancer cell CCTs (white arrows) invading into compact GBM tumor tissues composed by incomplete GBM cancer cells. There are multiple GBM cancer cell CCTs aggressively invade and extend into neighbor highly compacted GBM tumor tissues. GBM cancer cells in migration in CCT (purple arrows) are shown. Scale bar, 10μm.

**Fig. S6. Glioblastoma (GBM) cancer cells migrate in GBM CCTs and CCT networks, and disseminate and metastasize into far distance destinations in/beyond the brain.**

(A) Representative fluorescence image of GBM CCTs and CCT networks in cerebellum in clinical GBM tumors. There are many long, thin GBM CCTs degraded into CCT strands (CTS, orange arrows). Most of GBM cancer cells have left via dissemination in GBM CCTs and CCT networks, and the GBM cancer cell density in these areas are low, comparing to the highly compact and dense GBM cancer cells in the same GBM tissue in the lower part. Scale bar, 10μm.

(B) Representative fluorescence image of degradation of GBM CCTs and CCT networks in the brain. GBM CCTs degrade into countless GBM CCT strands (CTS, orange arrows) with variable sizes and in highly curved, coiled and irregular shapes, and form GBM CCT strand masses. Most (>99%) GBM cancer cells left this area via GBM CCTs and CCT networks and only few GBM cancer cells remain. There are a few GBM CCT cavities (white arrows) caused by GBM CCT degradation, decomposition and disappearance. Scale bar, 10μm.

**Fig. S7. Glioblastoma (GBM) cancer cell CCT degradation, CCT network decay, CNTS decomposition, micro blood vessel breakage and red blood cells spreading in the brain tissues.**

(A) Representative fluorescence image of clinical GBM tissue in the late stage. All GBM cancer cell CCTs are degraded into very thin CCT strand (CTS, orange arrows), and form cloud-like masses. Most GBM cancer cells left to neighbor and/or far-distance destinations via GBM CCTs and networks, and only a few GBM cancer cells remain. Some micro blood vessels are broken and leaky due to the process that many GBM CCTs invade into micro blood vessels, and many red blood cells (cyan arrows) spread and are randomly distributed in the brain tissues with significantly reduced cell density. Scale bar, 10μm.

(B) Representative fluorescence image of clinical GBM tissue in the very late stage. All GBM cancer cell CCTs are decomposed and most GBM CCT strands (CTS, orange arrows) disappeared. Micro blood vessels in this area are decomposed, and red blood cells (cyan arrows) randomly spread and distribute in the tumor liquefaction brain tissues. Scale bar, 10μm.

**Fig. S8. Cytocapsular cancer evolution in pancreas cancers.**

Representative fluorescence image of clinical pancreas cancer with a few CCTs (or CCT strands) (**A**), large quantities of CCTs, CCT networks (CCT strands) (**B**), and CCT degradation, CNTS decomposition (**C**). CCT (white arrows), CCT strand (CTS, orange arrows) are shown. Scale bar, 10μm.

**Fig. S9. Cytocapsular cancer evolution in thyroid cancers.**

Representative fluorescence image of clinical thyroid cancer with a few CCTs (or CCT strands) (**A**), large quantities of CCTs, CCT networks (CCT strands) (**B**), and CCT degradation, CNTS decomposition (**C**). CCT (white arrows), CCT strand (CTS, orange arrows) are shown. Scale bar, 10μm.

**Fig. S10. Cytocapsular cancer evolution in oral cancers.**

Representative fluorescence image of clinical oral cancer with a few CCTs (or CCT strands) (**A**), large quantities of CCTs, CCT networks (CCT strands) and superstructures of CCTs (**B**), and CCT degradation, CNTS decomposition (**C**). CCT (white arrows), CCT strand (CTS, orange arrows) are shown. Scale bar, 10μm.

**Fig. S11. Cytocapsular cancer evolution in prostate cancers.**

Representative fluorescence image of clinical prostate cancer with many CCTs (or CCT strands) (**A**), large quantities of CCTs, CCT networks, and CCT strands (**B**), and CCT degradation, CNTS decomposition (**C**). CCT (white arrows), CCT strand (CTS, orange arrows) are shown. Scale bar, 10μm.

**Fig. S12. Cytocapsular cancer evolution in lung cancers.**

Representative fluorescence image of clinical lung cancer with a few CCTs (or CCT strands) (**A**), many CCTs, CCT networks (CCT strands) (**B**), and large quantities of CCTs, CCT degradation, CNTS decomposition (**C**). CCT (white arrows), CCT strand (CTS, orange arrows) are shown. Scale bar, 10μm.

**Fig. S13. Cytocapsular cancer evolution in bone cancers.**

Representative fluorescence image of clinical bone cancer with a few bone colonies with CC (**A**), and large quantities of CCTs, CCT degradation, CNTS decomposition (**B**), CCTs, CCT networks, and many CCT degraded strands (**C**). CCT (white arrows), CCT strand (CTS, orange arrows) are shown. Scale bar, 10μm.

**Fig. S14. Cytocapsular cancer evolution in smooth muscle cancers.**

Representative fluorescence image of clinical smooth muscle cancer with a few CCTs (or CCT strands) (**A**), more CCTs, and CCT strands (**B**), and large quantities of CCTs, CCT degradation, CNTS decomposition (**C**). CCT (white arrows), CCT strand (CTS, orange arrows) are shown. Scale bar, 10μm.

**Fig. S15. Cytocapsular cancer evolution in fibrous tissue cancers.**

Representative fluorescence image of clinical fibrous tissue cancer with a few CCTs (or CCT strands) (**A**), more CCTs, and CCT strands (**B**), and large quantities of CCTs, CCT degradation, CNTS decomposition (**C**). CCT (white arrows), CCT strand (CTS, orange arrows) are shown. Scale bar, 10μm.

**Fig. S16. Cytocapsular cancer evolution in plasma cell myeloma in bone marrow. Representative** fluorescence image of clinical plasma cell myeloma in bone marrow with highly dense plasma cell myeloma, without CC/CCTs (or CCT strands) (**A**), a few CCTs and CCT strands (**B**), and large quantities of CCTs, CCT degradation, CNTS decomposition (**C**). CCT (white arrows), CCT strand (CTS, orange arrows) are shown. Scale bar, 10μm.

**Table S1. Cytocapsular cancer evolution analyses of 311 kinds/subtypes of cancers.**

## Notes

### Competing Interest Statement

The authors have declared no competing interest.

