## Supplementary material for "Cytocapsular cancer evolution analyses of 311 kinds of cancers": supple file

Fig. S1

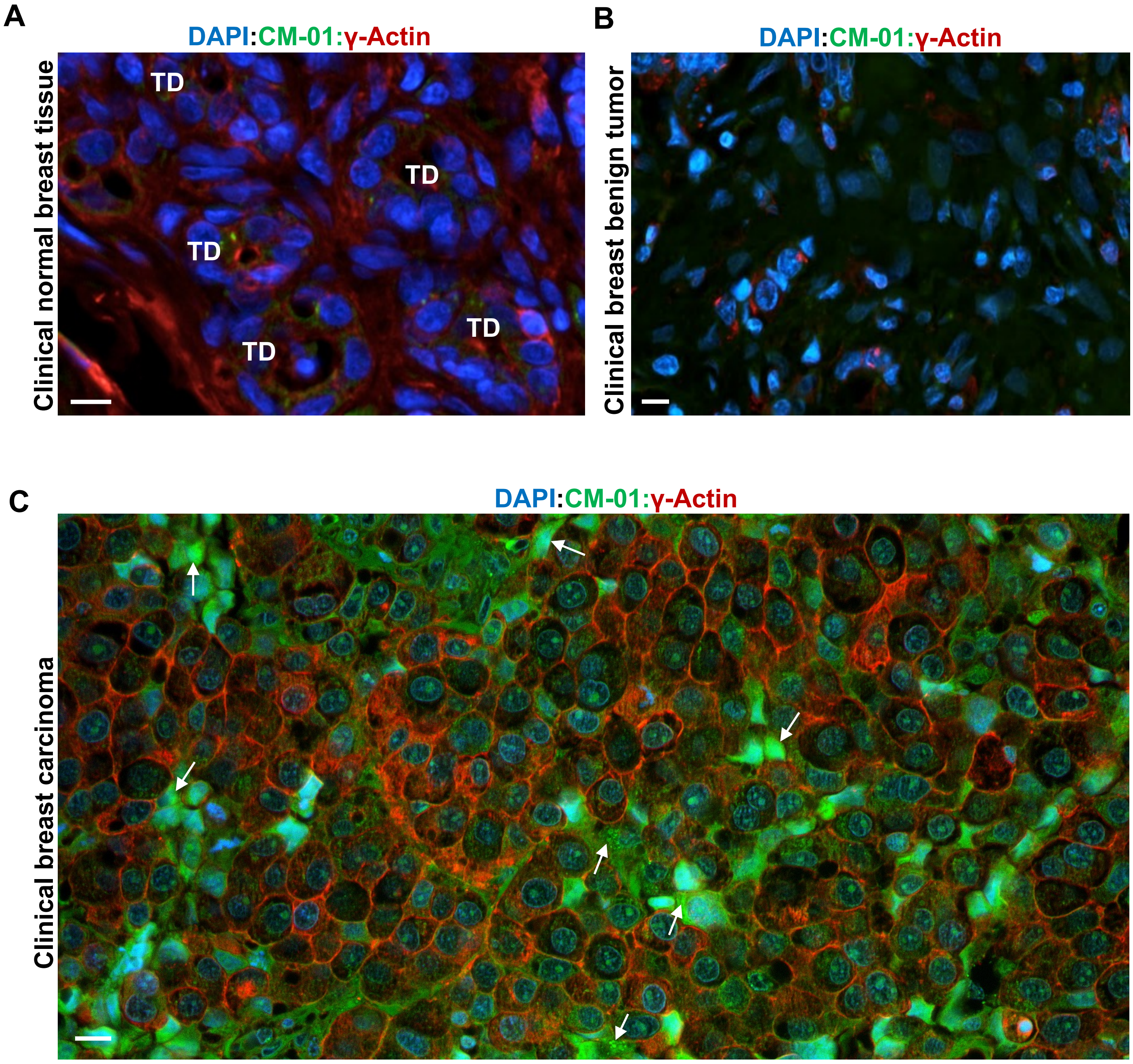
A
DAPI:CM-01:γ-Actin
Clinical normal breast tissue
TD
TD
TD
TD
TD
B
DAPI:CM-01:γ-Actin
Clinical breast benign tumor
C
DAPI:CM-01:γ-Actin
Clinical breast carcinoma

Fig. S2

Breast-Invasive Ductal Carcinoma

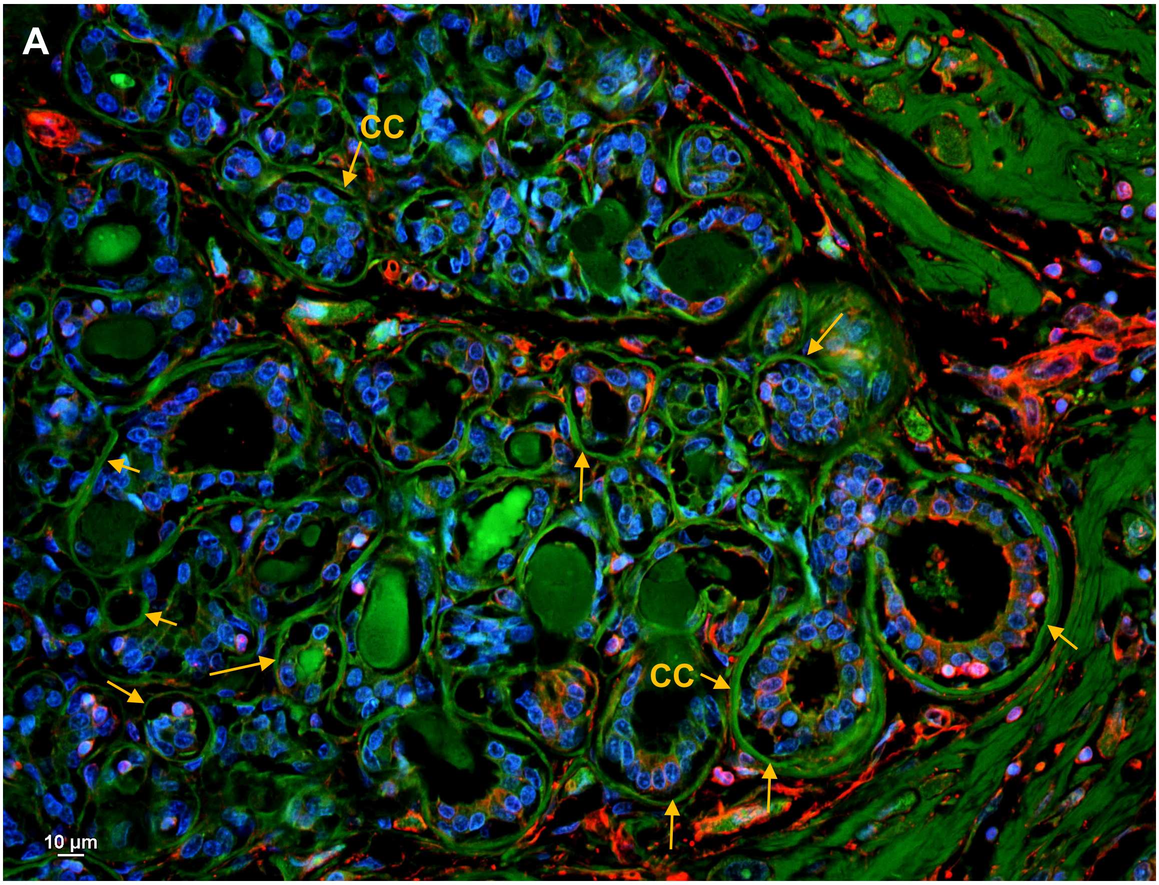


DAPI:CM-01:γ-Actin

Breast-Invasive Ductal Carcinoma

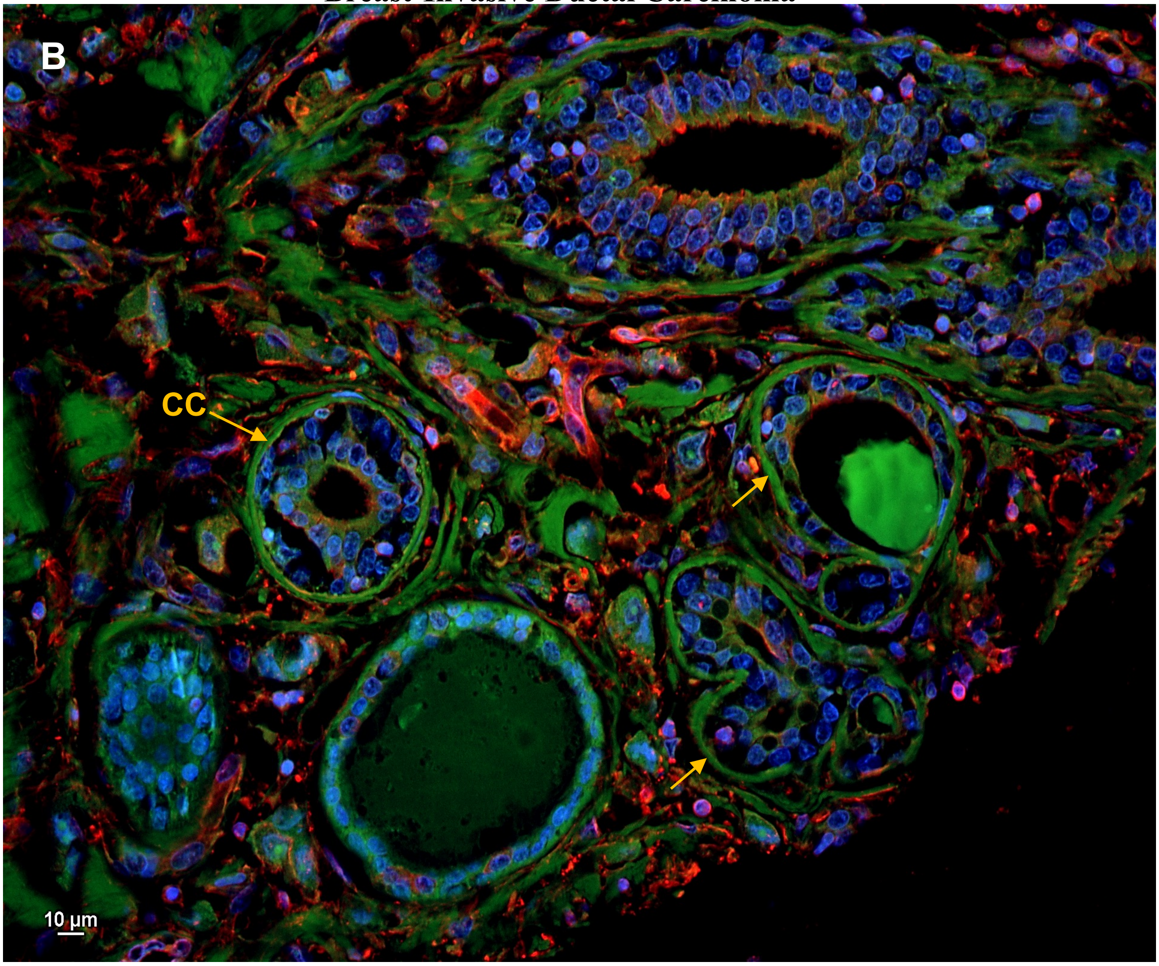


DAPI:CM-01:γ-Actin

Fig. S3

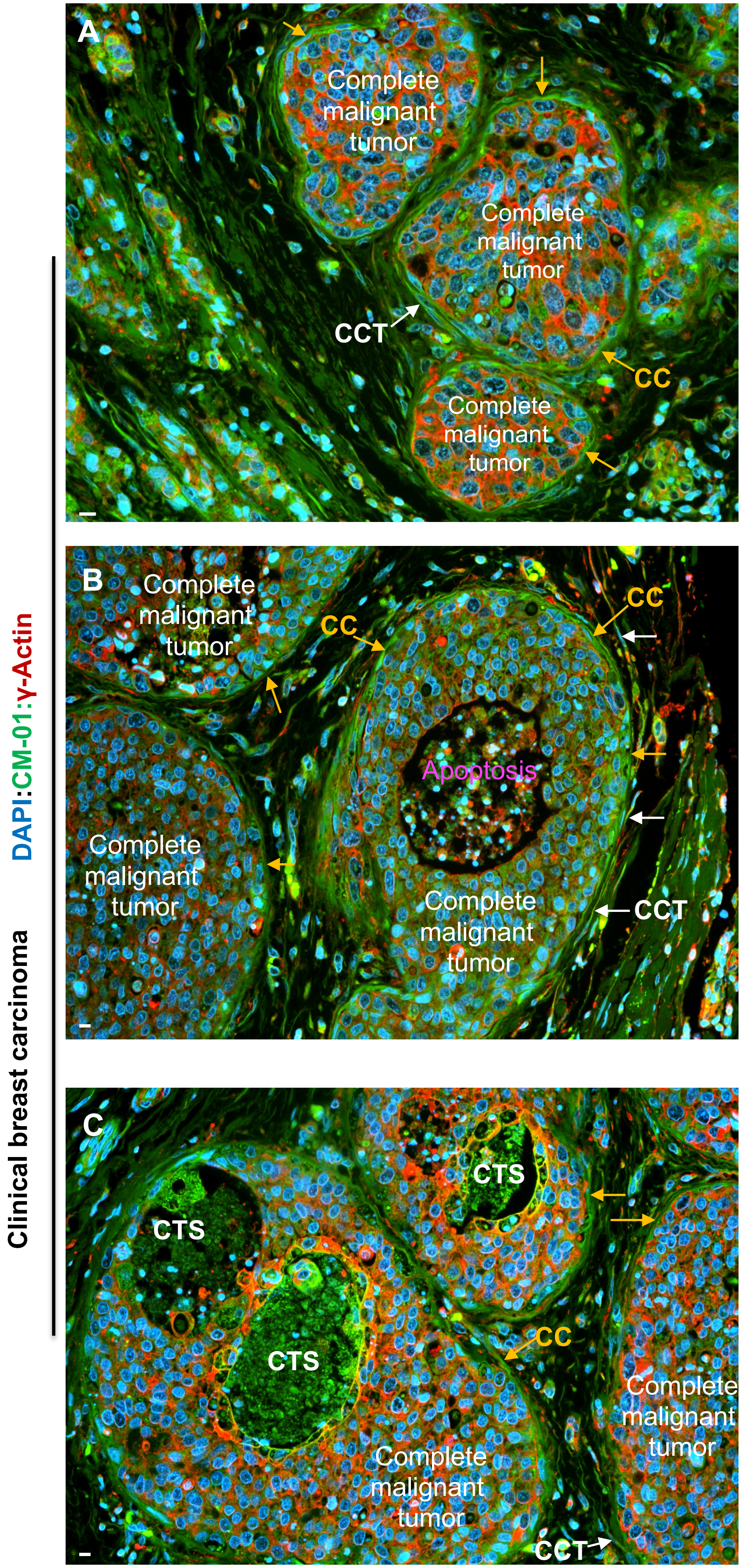

A
Complete malignant tumor
Complete malignant tumor
Complete malignant tumor
CCT
CC
B
Complete malignant tumor
CC
CC
Apoptosis
Complete malignant tumor
Complete malignant tumor
CCT
C
CTS
CTS
CTS
CC
Complete malignant tumor
Complete malignant tumor
CCT
DAPI:CM-01:γ-Actin
Clinical breast carcinoma

**Fig. S4**

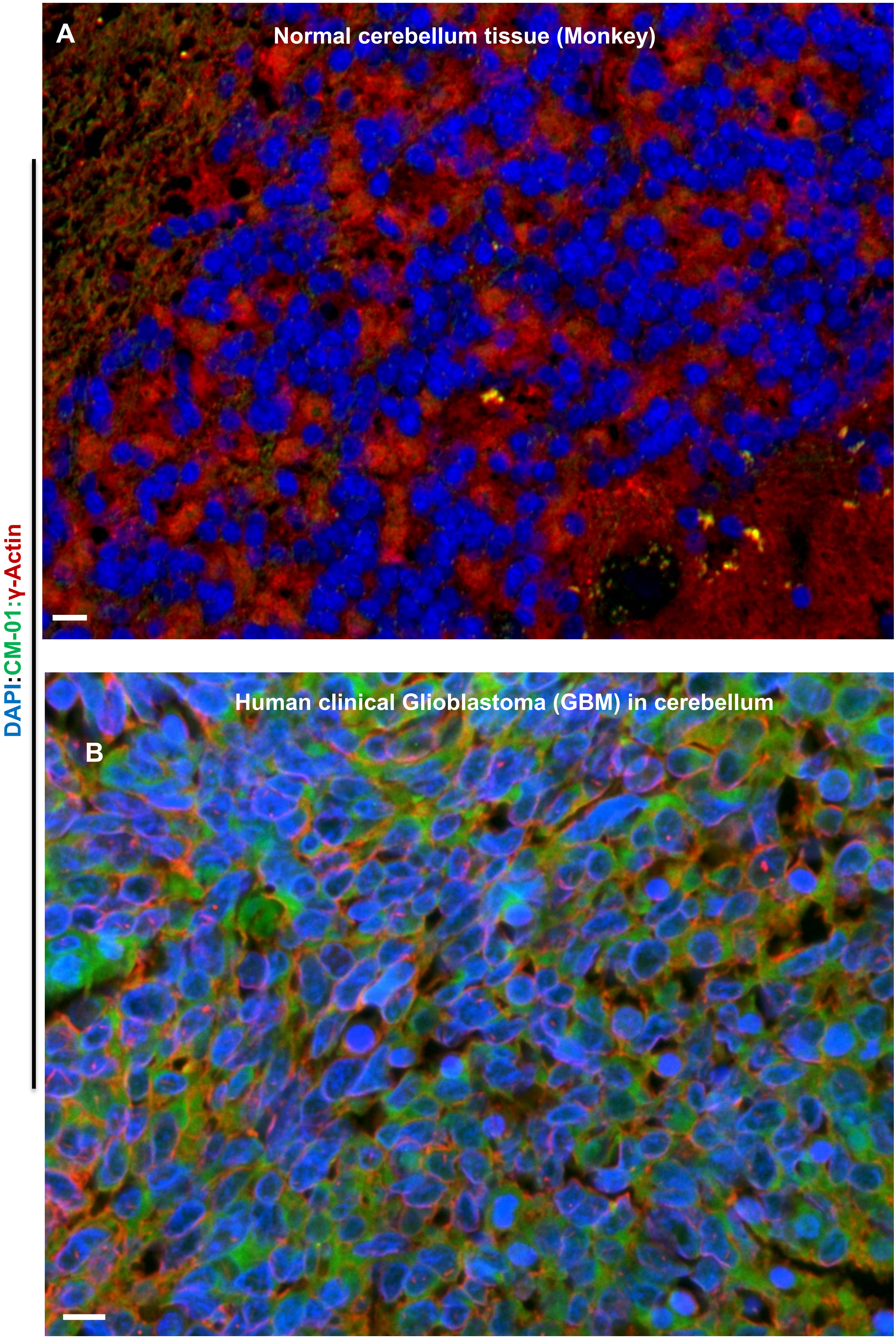

A
Normal cerebellum tissue (Monkey)
B
Human clinical Glioblastoma (GBM) in cerebellum
DAPI:CM-01:γ-Actin

Fig. S5

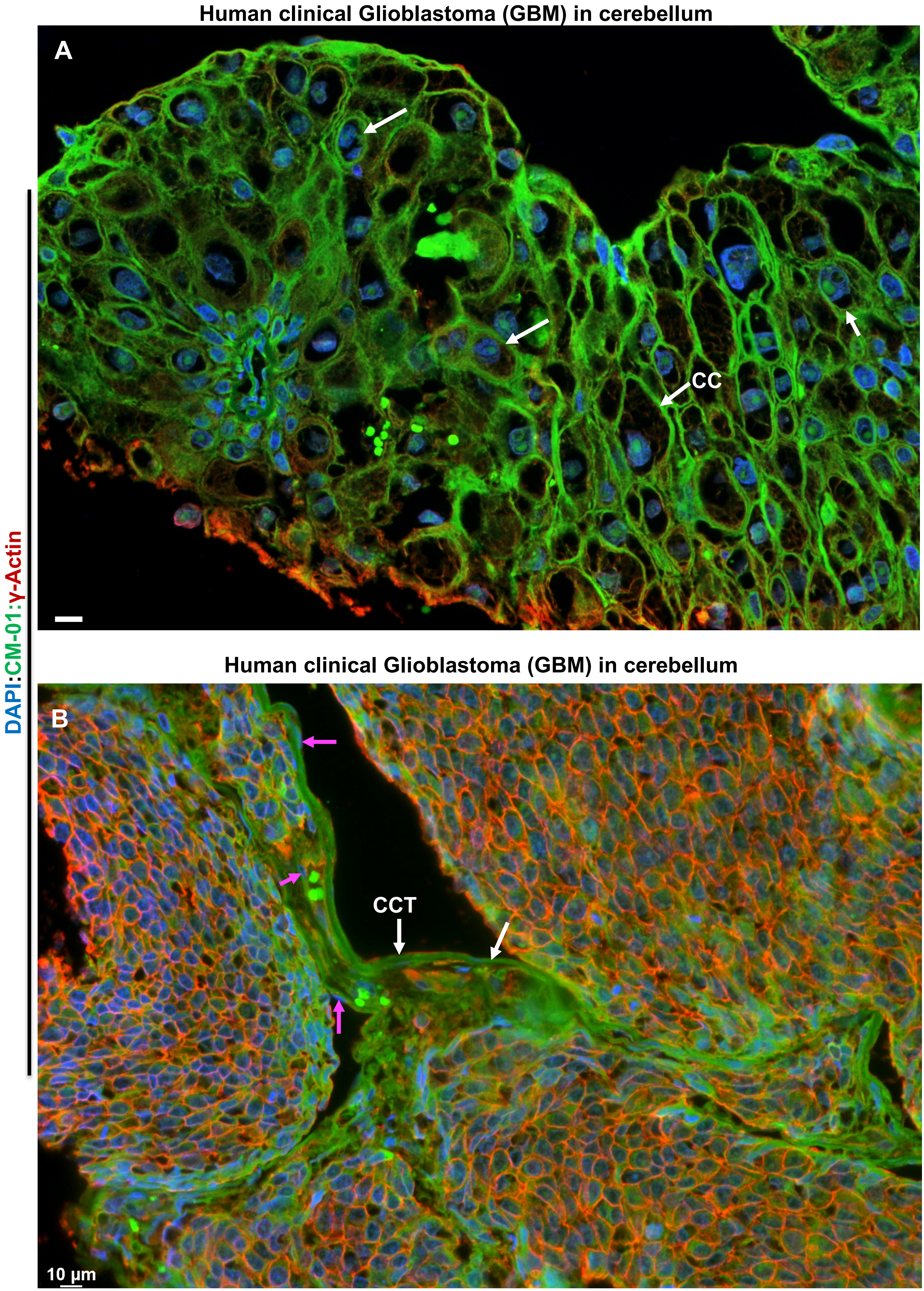
Human clinical Glioblastoma (GBM) in cerebellum
A
CC
Human clinical Glioblastoma (GBM) in cerebellum
B
CCT
10 µm
DAPI:CM-01:γ-Actin

Fig. S6

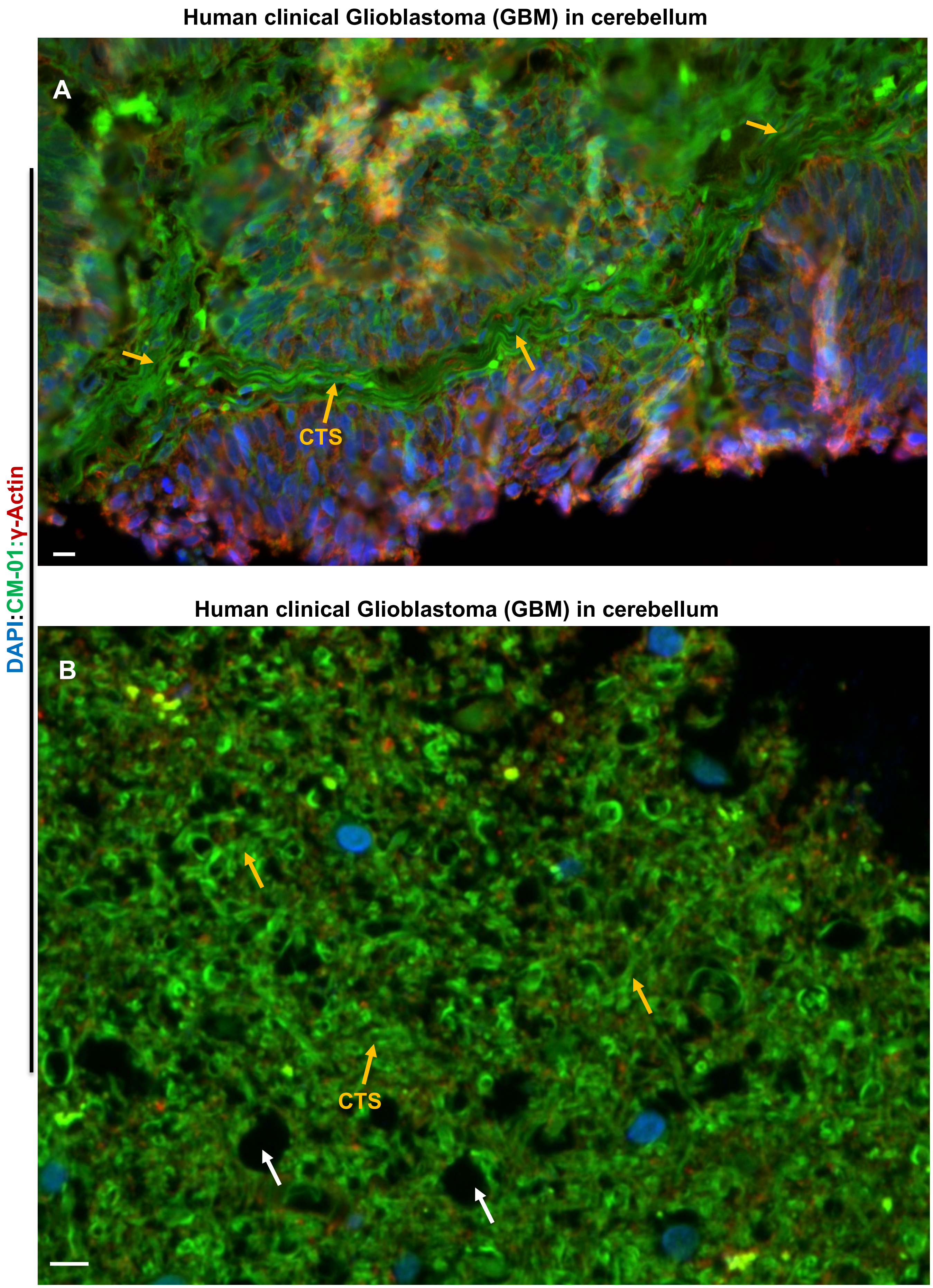
Human clinical Glioblastoma (GBM) in cerebellum
A
CTS
Human clinical Glioblastoma (GBM) in cerebellum
B
CTS
DAPI:CM-01:γ-Actin

**Fig. S7**

**Human clinical Glioblastoma (GBM) in cerebellum**

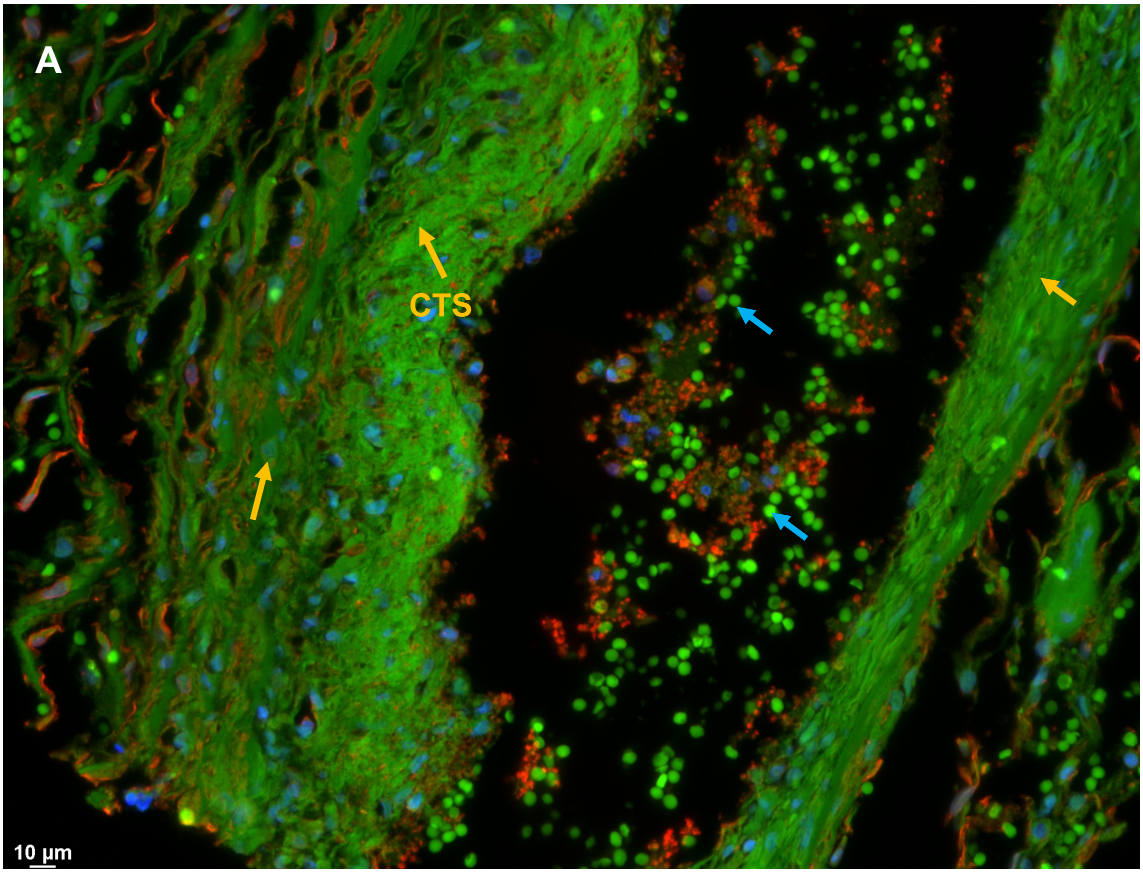


**Human clinical Glioblastoma (GBM) in cerebellum**

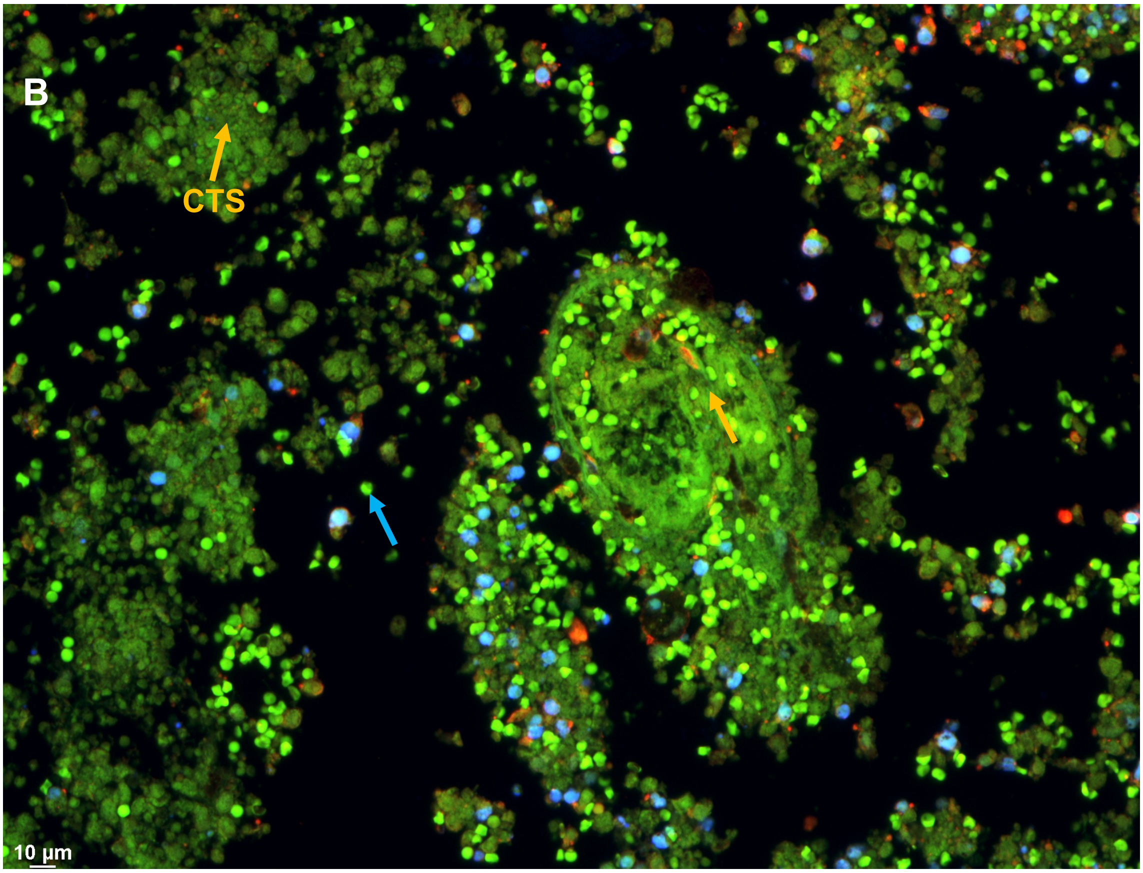

Fig. S8

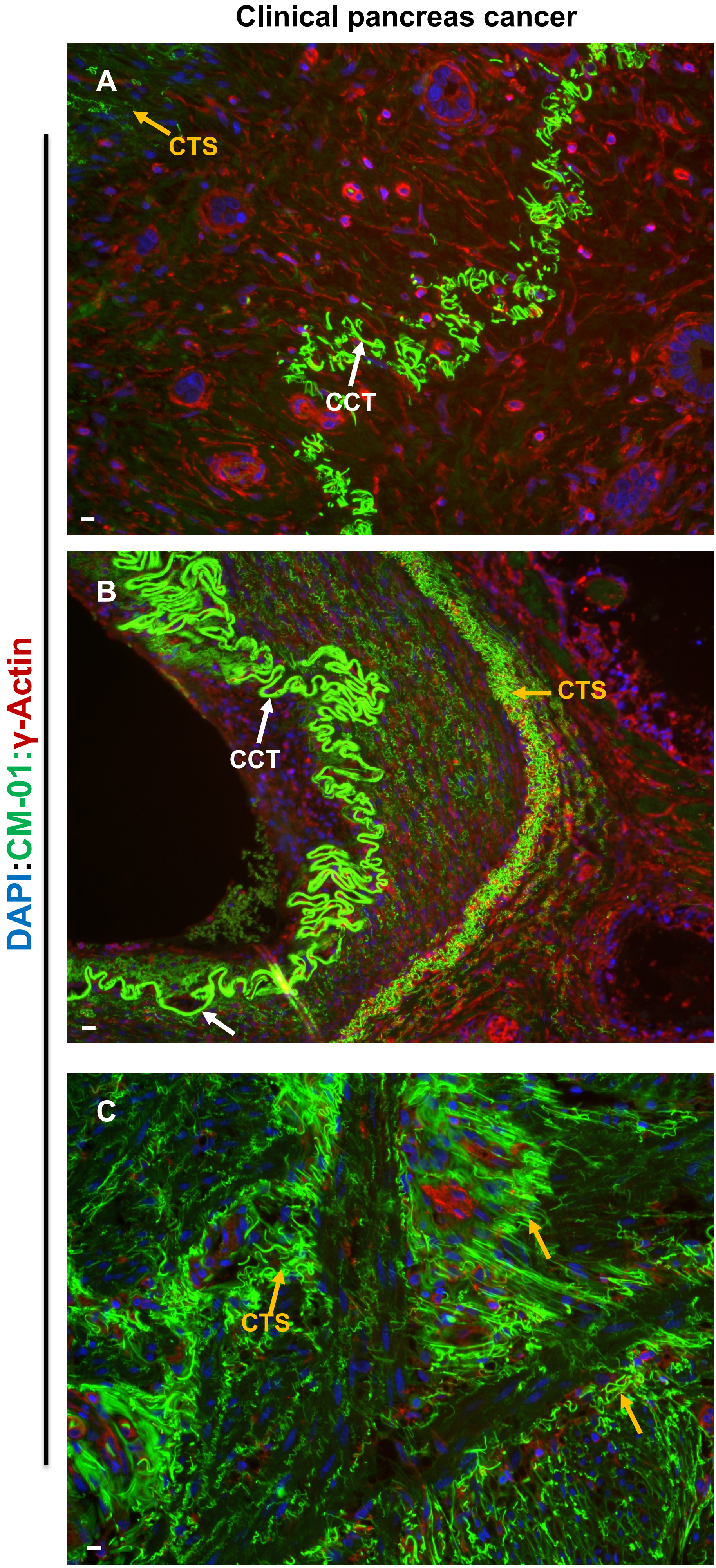
Clinical pancreas cancer
DAPI:CM-01:γ-Actin
A
CTS
CCT
B
CTS
CCT
C
CTS

**Fig. S9**

### Clinical thyroid cancer

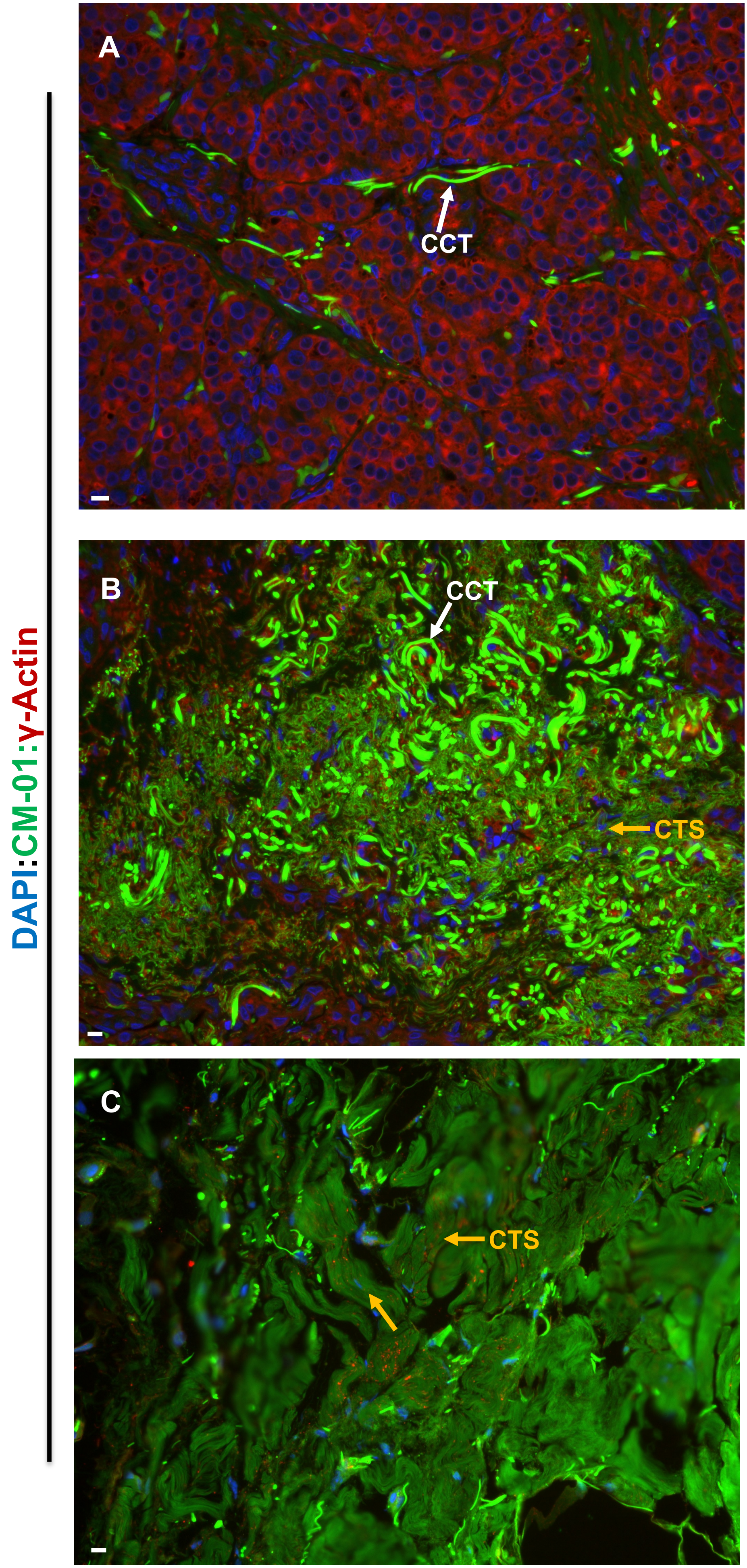

Fig. S10

Clinical oral cancer

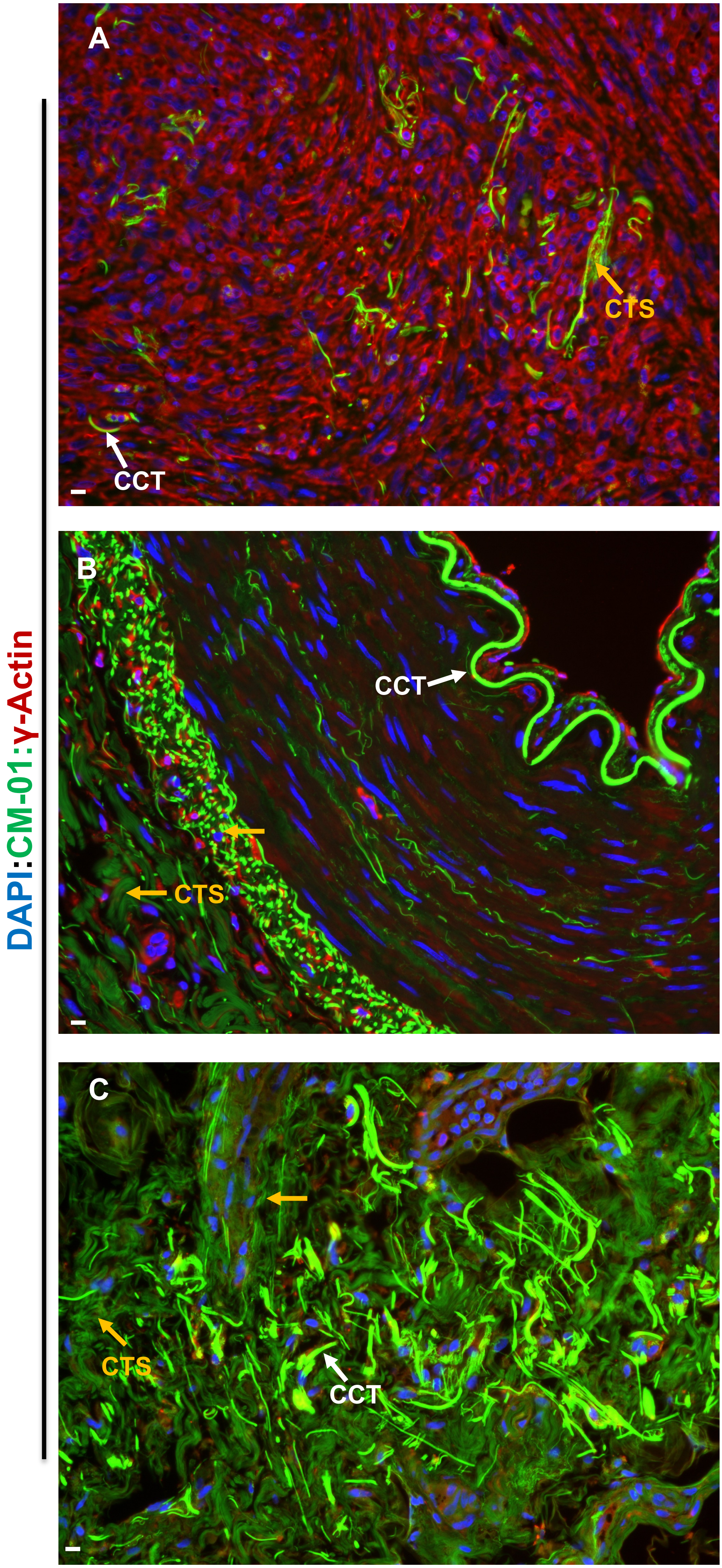

Fig. S11

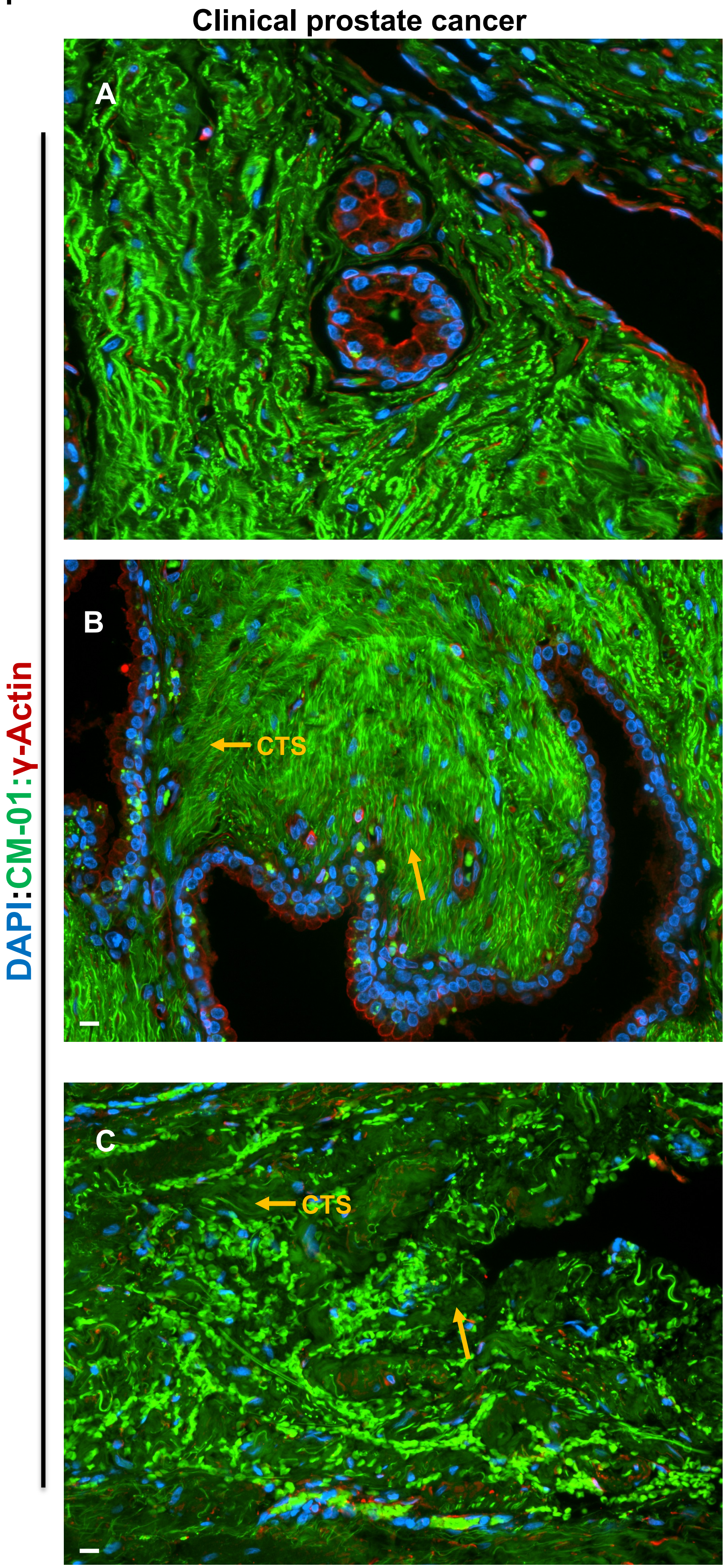

Fig. S12

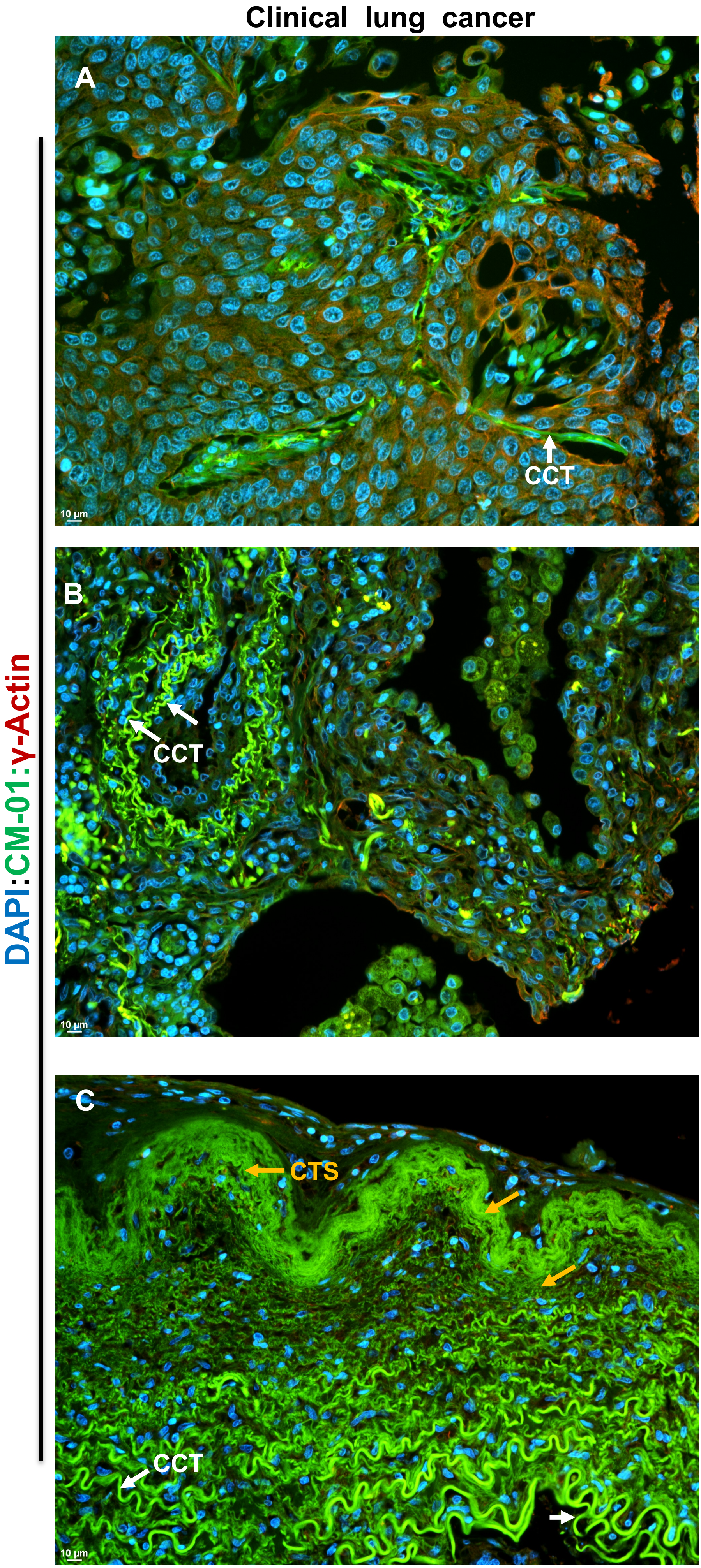
Clinical lung cancer
DAPI:CM-01:γ-Actin
A
CCT
10 µm
B
CCT
10 µm
C
CTS
CCT
10 µm

Fig. S13

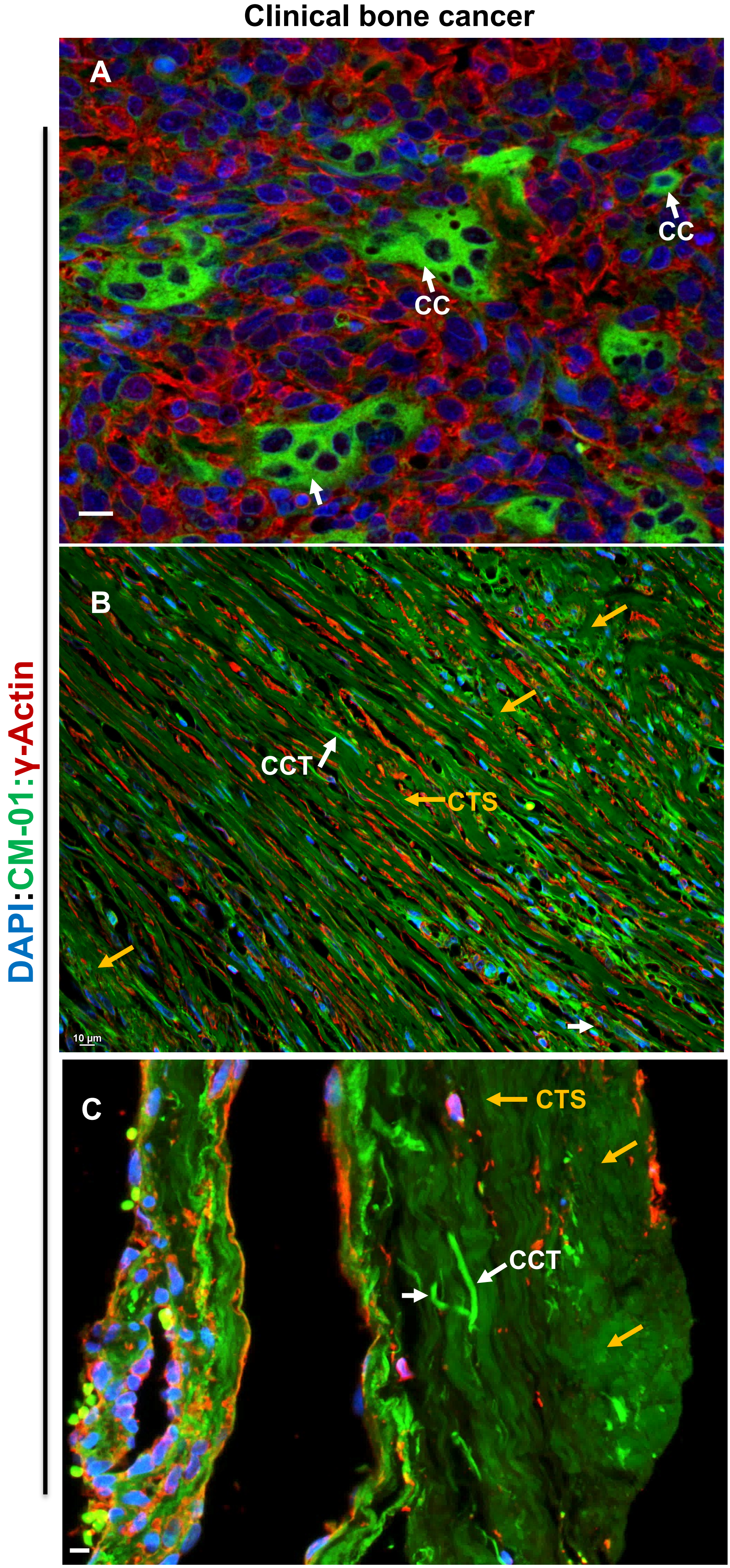

Clinical bone cancer
A
CC
CC
B
CCT
CTS
10 µm
C
CTS
CCT
DAPI:CM-01:γ-Actin

Fig. S14

Clinical smooth muscle cancer

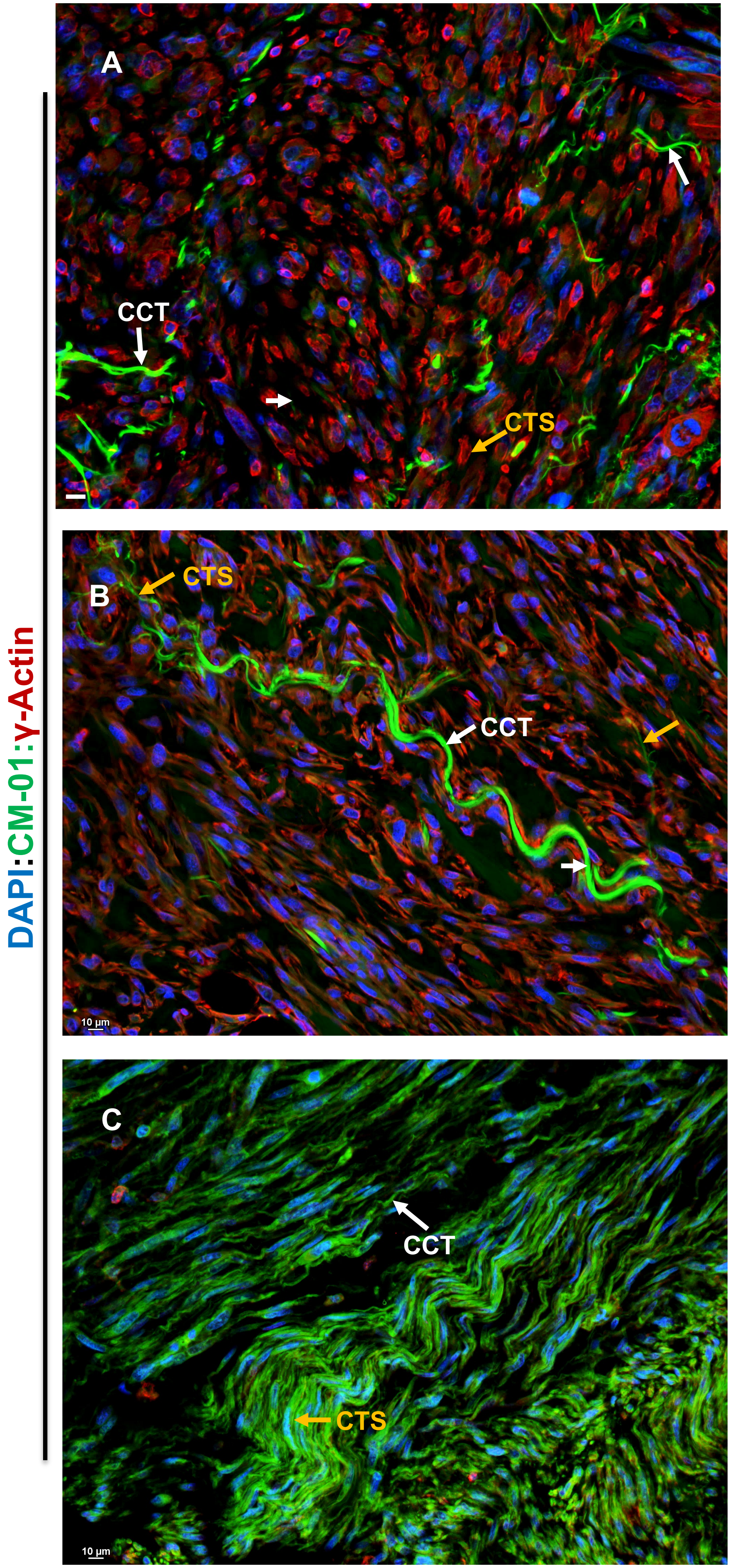

Fig. S15

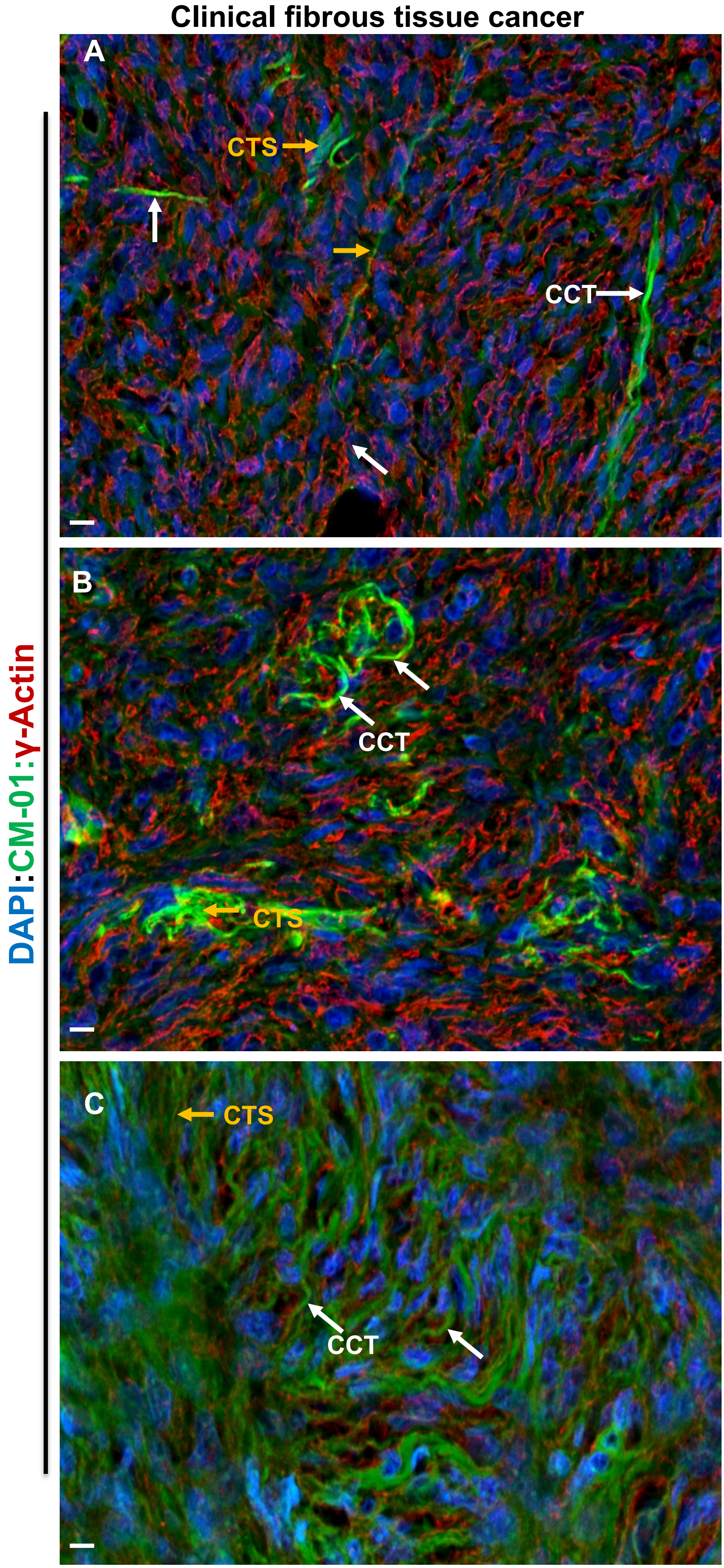

Clinical fibrous tissue cancer
A
CTS
CCT
B
CCT
CTS
C
CTS
CCT
DAPI:CM-01:γ-Actin

Fig. S16

Clinical plasma cell myeloma in bone marrow

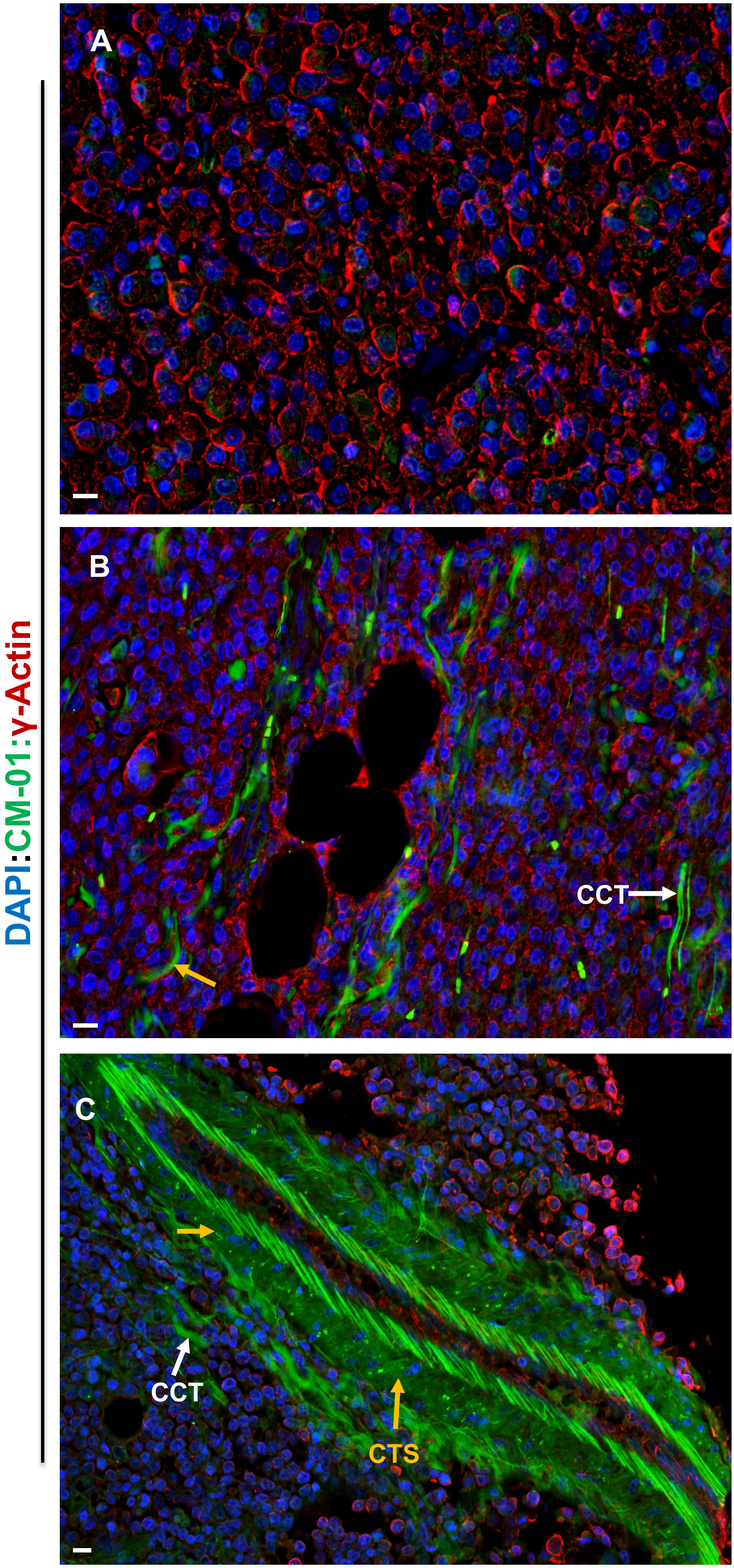

**Table S1.** Cytocapsular cancer evolution analyses of 311 kinds/subtypes of cancers.

| | Cancer types by organs/ tissues | Cancer subtypes | Tissue specimen number | Patient number | Male | Female | Age | Cancer stages | With incomplete cancer cells | Complete cancer cells with CCs | With CCT initiation | With CCTs | With complete tumors in CC | With CCT network | With CNTS | With moving CNTS |
|---|---|---|---|---|---|---|---|---|---|---|---|---|---|---|---|---|
| 1 | Adrenal Gland | 3 | 23 | 22 | 10 | 12 | 26-71 | I-IV | + | + | + | + | + | + | + | + |
| 2 | Bladder | 7 | 46 | 46 | 30 | 16 | 36-74 | I-IV | + | + | + | + | + | + | + | + |
| 3 | Blood (in blood vessel) | 9 | 14 | 14 | 8 | 6 | 21-45 | I-IV | + (cc)# | + (cc)# | | | | | | |
| 4 | Blood (in bone marrow) | 28* | 35 | 35 | 18 | 17 | 35-70 | I-IV | + | + | + | + | + | + | + | + |
| 5 | Bone | 12 | 212 | 210 | 115 | 95 | 21-78 | I-IV | + | + | + | + | + | + | + | + |
| 6 | Bone marrow | 6 | 20 | 20 | 10 | 10 | 25-64 | I-IV | + | + | + | + | + | + | + | + |
| 7 | Brain | 3 | 124 | 122 | 86 | 36 | 24-67 | I-IV | + | + | + | + | + | + | + | + |
| 8 | Breast | 46 | 3960 | 3890 | | 3890 | 18-86 | In situ | + | + | + | + | + | + | + | + |
| | | | | | | | | I | + | + | + | + | + | + | + | + |
| | | | | | | | | II | + | + | + | + | + | + | + | + |
| | | | | | | | | III | + | + | + | + | + | + | + | + |
| | | | | | | | | IV | + | + | + | + | + | + | + | + |
| 9 | Cervix uteri | 7 | 53 | 51 | | 51 | 37-55 | I-IV | + | + | + | + | + | + | + | + |
| 10 | Colon | 6 | 1586 | 1586 | 823 | 763 | 16-90 | I-IV | + | + | + | + | + | + | + | + |
| 11 | Esophagus | 5 | 63 | 62 | 41 | 21 | 24-65 | I-IV | + | + | + | + | + | + | + | + |
| 12 | Fibrous | 2 | 25 | 25 | 12 | 13 | 35-67 | I-IV | + | + | + | + | + | + | + | + |
| 13 | Gallbladder | 3 | 26 | 26 | 20 | 6 | 37-62 | I-IV | + | + | + | + | + | + | + | + |
| 14 | Head/neck | 5 | 12 | 12 | 4 | 8 | 41-66 | I-IV | + | + | + | + | + | + | + | + |
| 15 | Intestine | 3 | 15 | 15 | 10 | 5 | 36-68 | I-IV | + | + | + | + | + | + | + | + |
| 16 | Kidney | 10 | 123 | 120 | 81 | 39 | 33-65 | I-IV | + | + | + | + | + | + | + | + |
| 17 | Liver | 6 | 362 | 358 | 231 | 127 | 36-76 | I-IV | + | + | + | + | + | + | + | + |
| 18 | Lung | 23 | 683 | 676 | 437 | 239 | 27-78 | I-IV | + | + | + | + | + | + | + | + |
| 19 | Lymph | 9 | 56 | 56 | 31 | 25 | 21-73 | I-IV | + | + | + | + | + | + | + | + |
| 20 | Oesophagus | 2 | 4 | 4 | 3 | 1 | 34-54 | I-IV | + | + | + | + | + | + | + | + |
| 21 | Oral cavity | 5 | 17 | 17 | 10 | 7 | 28-69 | I-IV | + | + | + | + | + | + | + | + |
| 22 | Ovary | 7 | 83 | 83 | | 83 | 32-61 | I-IV | + | + | + | + | + | + | + | + |
| 23 | Pancreas | 13 | 222 | 218 | 162 | 56 | 28-74 | I-IV | + | + | + | + | + | + | + | + |
| 24 | Penis | 2 | 3 | 3 | 3 | | 45-62 | I-IV | + | + | + | + | + | + | + | + |
| 25 | Prostate | 25 | 1606 | 1528 | 1528 | | 35-77 | I-IV | + | + | + | + | + | + | + | + |
| 26 | Rectum | 5 | 5 | 5 | 4 | 1 | 32-57 | I-IV | + | + | + | + | + | + | + | + |
| 27 | Skin | 7 | 122 | 122 | 35 | 87 | 21-79 | I-IV | + | + | + | + | + | + | + | + |
| 28 | Spleen | 2 | 6 | 6 | 1 | 5 | 36-48 | I-IV | + | + | + | + | + | + | + | + |
| 29 | Smooth muscle | 2 | 20 | 20 | 10 | 10 | 35-55 | I-IV | + | + | + | + | + | + | + | + |
| 30 | Stomach | 26 | 245 | 245 | 123 | 122 | 36-72 | I-IV | + | + | + | + | + | + | + | + |
| 31 | Testis | 5 | 24 | 24 | 24 | | 41-67 | I-IV | + | + | + | + | + | + | + | + |
| 32 | Thymus | 4 | 5 | 5 | 4 | 1 | 37-56 | I-IV | + | + | + | + | + | + | + | + |
| 33 | Thyroid | 5 | 17 | 17 | 10 | 7 | 29-61 | I-IV | + | + | + | + | + | + | + | + |
| 34 | Uterus | 5 | 26 | 26 | | 26 | 35-56 | I-IV | + | + | + | + | + | + | + | + |
| 35 | Vulva | 3 | 13 | 13 | | 13 | 32-55 | I-IV | + | + | + | + | + | + | + | + |
| | Total | 311 | 9856 | 9682 | 4227 | 5455 | | | | | | | | | | |
